# An ancient lysozyme in placozoans participates in acidic extracellular digestion

**DOI:** 10.1101/2024.10.06.616844

**Authors:** Henry Berndt, Igor Duarte, Urska Repnik, Michel Struwe, Mohammad Abukhalaf, Axel Scheidig, Andreas Tholey, Harald Gruber-Vodicka, Matthias Leippe

**Author notes:** Corresponding authors e-mail addresses –.

## Abstract

Lysozymes are an essential part of immunity and nutrition in metazoans, degrading bacterial cell walls via the hydrolysis of peptidoglycan. Although various lysozymes have been reported for higher animals, the origin of animal lysozymes remains elusive as they seem to be lacking in all early branching phyla. In this study, we investigated a putative goose-type lysozyme (PLys, glycoside hydrolase family 23, GH23) of the placozoan Trichoplax sp. H2. We show that PLys is highly active and produced in gland cells of the ventral epithelium. PLys contains a protective and non-conserved cysteine-rich domain N-terminal of the conserved GH23 lysozyme domain. A truncation of this N-terminal domain in the maturation process of PLys leads to a drastic increase in enzymatic activity at the cost of stability. As the lysozyme is most active under acidic conditions, we investigated the pH trajectories during extracellular digestion *in situ*. Using a pH-senstive fluorescence reporter, we show that *Trichoplax* sp. H2 acidifies its temporary feeding grooves pulsatively during digestive events close to the optimum pH for PLys activity. To elucidate the evolutionary origin of the metazoan GH23 lysozyme family, we applied a structure-based phylogenetics approach to show that the metazoan g-type GH23 lysozymes originated from a horizontal gene transfer event from bacteria to an early pre-bilaterian ancestor. GH23 lysozymes have then been retained and expanded in many phyla, including Porifera, Cnidaria, Placozoa and chordates, acting as first animal lysozyme and a key component in the antibacterial arsenal since early animal evolution.

## Main

Animals are always associated and interacting with bacteria. These interactions can range from beneficial mutualism between host and microbe to one side taking advantage, e. g. bacterial pathogens or bacterivorous animals (1). For killing and digestion of bacteria, animals feature a variety of molecular processes and effector proteins. A key group of such antibacterial proteins are lysozymes, glycoside hydrolases (GHs) that attack the peptidoglycan layer by cleaving the β-1,4-glycosidic bond between its main building blocks N-acetylmuramic acid (MurNAc) and N-acetylglucosamine (GlcNAc). Lysozymes have been traditionally grouped into different classes, named after the organisms in which they were first discovered (2). A more consistent naming convention for enzymes with a glycoside hydrolase activity categorizes them into families based on their fold and substrate specificity. For example, goose-type (g-type) lysozymes, first discovered in the egg white of the graylag goose *Anser anser* and found later in many vertebrates, belong to the glycoside hydrolase family 23 (GH23), together with bacterial lytic transglycosylases (3).

Lysozymes appear to be very unevenly distributed in the animal tree of life, with most early branching clades such as Ctenophora and Cnidaria apparently lacking *bona fide* lysozymes (2). While protostomian invertebrates as well as early branching deuterostomes such as echinoderms predominantly feature invertebrate-type (i-type) lysozymes of GH22i, chordates primarily possess g-type (GH23) and chicken-type (c-type) lysozymes (GH22c) (2,4). Sparse reports of c-type lysozymes in Arthropoda and g-type lysozymes in mollusks, however, challenge the evolutionary argument that these two lysozyme families are limited to Chordata (2). In addition, a short report about a putative g-type in placozoans, provisionally named PLys, suggests that the evolution of major lysozyme groups in animals was not a relatively late innovation in the lineages leading to bilaterians and then chordates (5). The existence of a potential g-type lysozyme in some of arguably the simplest metazoan animals points instead to a much earlier occurrence and diversification of lysozymes in the animal tree of life (ATL). The position of Placozoa in the ATL is under ongoing discussion, but recent bioinformatic analyses consistently place placozoans among the pre-bilaterian lineages and as the sister taxon to the clade formed by Cnidaria and Bilateria (6,7). With such an interesting position in animal phylogeny, placozoans are increasingly used as a model organism to study the origin of multicellularity, developmental biology, and neurophysiology (8–10). As millimeter-sized and disk-shaped grazers of marine biofilms, placozoans continually interact with bacteria, and also harbor endosymbiotic bacteria in a cell-type specific symbiosis (11,12). The basic body plan of the placozoans is very simple and consists of three layers. The ventral columnar epithelium has a mainly digestive function, whereas a thin dorsal epithelium has primarily a barrier function. A densely packed mesenchyme-like layer consists of fiber-cells, which form many cell-to-cell connections with other cell types in both epithelia (13). In addition, fiber cells contain endosymbiotic bacteria, and have been highlighted as central immune cells in placozoa, as they are the only cells capable of phagocytosis and apoptosis (11,14). During feeding, placozoa form a temporary gastric cavity between the ventral epithelium and the substrate, into which gland cells and lipophilic cells secrete digestive enzymes (15). Comparative single-cell transcriptomics data of diverse placozoan lineages showed *plys* to be almost exclusively transcribed in gland cells of the ventral epithelium, from where PLys could be secreted into the gastric cavity (6).

To elucidate the role of PLys in *Trichoplax* sp., we identified the natural proteoforms of the lysozyme, produced the two lysozyme variants recombinantly, characterized them *in vitro* and determined their cellular distribution in whole animals. Using evolutionary analyses, we show that the highly active lysozyme PLys represents a broad radiation of g-type lysozymes across the animal tree of life that arguably were among the earliest antibacterial effectors in metazoan evolution.

## Results and Discussion

### Identification and localization of native placozoan lysozyme

As a putative g-type lysozyme has been reported for placozoans (PLys) (5) and the corresponding orthologous gene is transcribed across different placozoan species (Fig. S1), we screened for the activity of lysozymes using placozoan total protein extract from *Trichoplax* sp. H2. On a lysoplate, placozoan total protein extract in two concentrations (0.24 mg/mL and 1.24 mg/mL) produced zones of lysis after 24 h (Fig. 1A). Following this first indication of lysozyme activity, we validated the identity of the potential lysozymes by mass spectrometry and Western blotting of a reversed-phase chromatography fractionated *Trichoplax* proteome. The Western blot revealed two protein double bands, which correspond to molecular masses of about 30 kDa and 20 kDa, matching the two proteoforms previously mentioned by Bam and Nakano (Fig. S2) (5). We therefore used fractions with the strongest signals in the western blot for mass spectrometry to characterize these two potential proteoforms in a combination of proteomic analyses of intact proteins without further processing (top-down) and of proteolytically generated protein fragments (peptide-based; bottom-up). In top-down experiments, the most abundant proteoform was lacking the N-terminus and started at amino acid residue S100 (Fig. 1B). Based on the structural model of the full length PLys predicted by ColabFold, the N-terminal domain that is missing in this most abundant proteoform consists of two repeated beta-sheets, is stabilized by 6 disulphide bonds, and is connected to the active g-type lysozyme domain by a loop. N-terminomics (Bottom-up proteomics) experiments detected peptides specific to this N-terminal domain of the full-length PLys. The N-terminus (S100) of the shorter form of PLys is located in the connective loop region between the N-terminal domain and the lysozyme domain (Fig. 1C), which likely constitutes a cleavage site for specific proteolytic processing during protein maturation (limited proteolysis). We therefore differentiate between the truncated and presumably mature PLys (mPLys) and the full-length precursor PLys (pPLys) in the later experiments.

**Figure 1:**
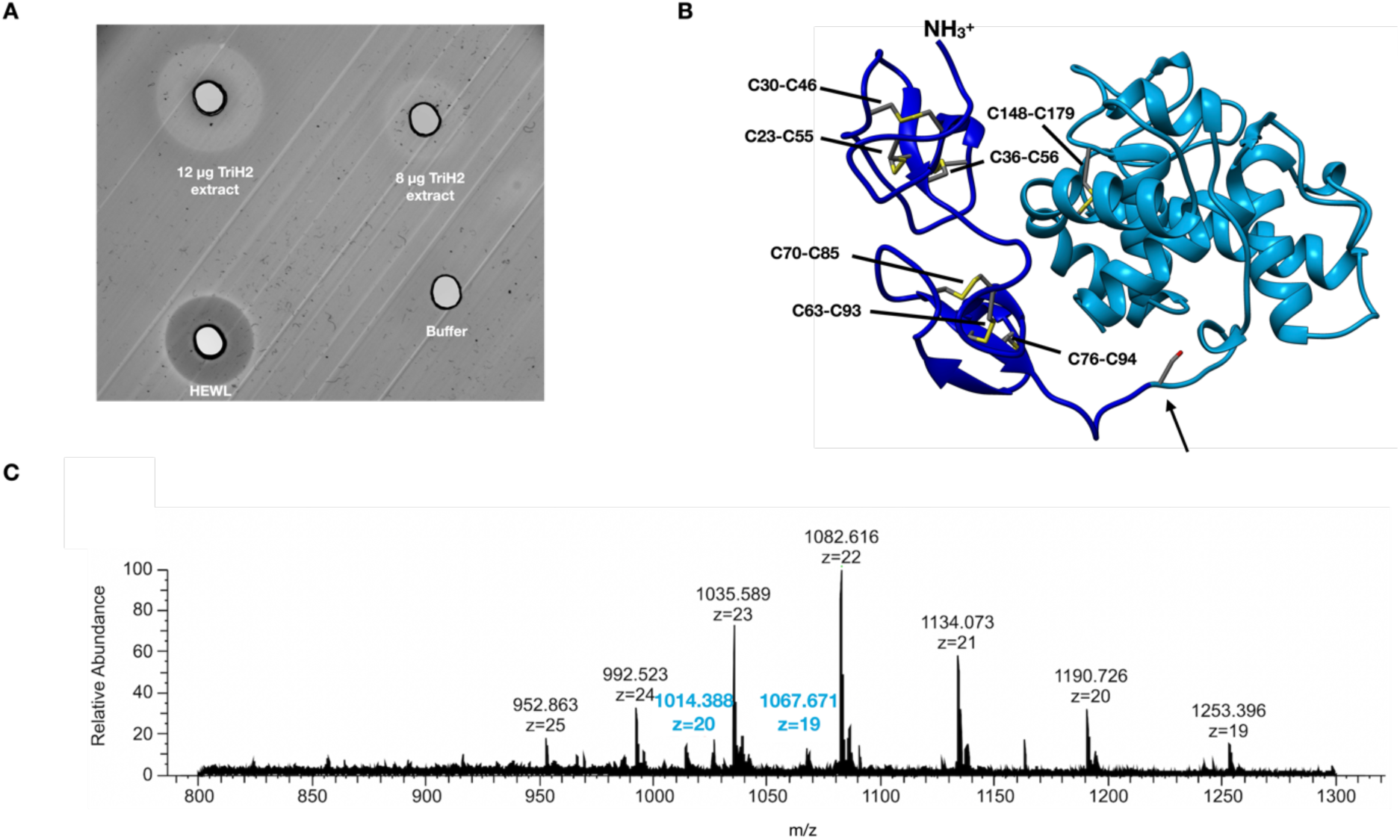
PLys is an active lysozyme with a truncated mature proteoform. **A**): Lysoplate assay with 8 μg and 12 μg total protein of raw *Trichoplax* sp. H2 protein extract, 20 μg hen egg white lysozyme (HEWL), and lysis buffer. **B)**: Structure model of PLys as ribbon diagram. The backbone of the mature proteoform of PLys is colored in light blue, while the cysteine-rich N-terminal domain is colored in dark blue. Cysteine residues forming disulphide bonds are highlighted and annotated. An arrow marks S100, the apparent N-terminus of the physiologically mature proteoform. **C**): Representative mass spectrum of the top-down proteomics experiment with the identified mature PLys proteoform with charge states z=19 and z=20.

Following the identification of the two forms of PLys in the *Trichoplax* sp. H2, we applied immunohistochemical labeling of PLys for cellular localization of the enzyme. Lysozymes are usually either secreted for extracellular lysis of bacteria or act on phagocytosed bacteria in phagolysosomes (2). As fiber cells have been indicated as the only cell type in *Trichoplax* capable of engulfing bacteria (14), we predicted that PLys is detectable either in lysosomes of fiber cells or in exocytotic vesicles of secretory cells of the ventral epithelium (gland cells or lipophilic cells). In general, placozoans feature an upper (dorsal) epithelium (d, Fig. 2A), made up by dorsal epithelial cells, and a lower (ventral) epithelium (v, Fig. 2A) which is populated by ventral epithelial cells, gland cells, and lipophilic cells (13,16). Lipophilic cells are characterized by large vesicles close to their apical membrane which are stainable using lipophilic and acidophilic dyes (15). A mucus secreting subtype of gland cells shows high abundance in the peripheral region of the ventral epithelium that is stainable with wheat germ agglutinin (WGA) (16). The region between the epithelia is mostly covered by fiber cells and to a smaller degree by gravity-sensing crystals cells (13,14). The signal of the PLys-specific polyclonal antibodies was present across the animal (Fig. 2B) and localized mostly to cells in the ventral epithelium (Fig. 2C). We also observed labeling in traces of mucus the animal left behind, which is another indicator for secretion of the enzyme (Fig. 2B). Additionally, we occasionally observed accumulation of signals between ventral and dorsal epithelium that could origin from fiber cells. Our antibody staining is consistent with single-cell expression data demonstrating *plys* expression predominantly in gland cells (6). Given the inconsistent anti-PLys-signal in fiber cells, it remains unclear, whether fiber cells are able to import gland-cell secreted lysozyme, or if the synthesis of lysozyme in fiber cells is inducible by bacterial components. While the genome of the closely related *Trichoplax adhaerens* appears to code for a high variety of scavenger receptors and reveals parts of Toll-like and NOD-like signaling pathways that may detect bacterial molecular patterns, complete signaling pathways from receptors to transcription factors that induce and regulate the synthesis of innate immunity factors such as PLys have yet to be identified (17).

**Figure 2:**
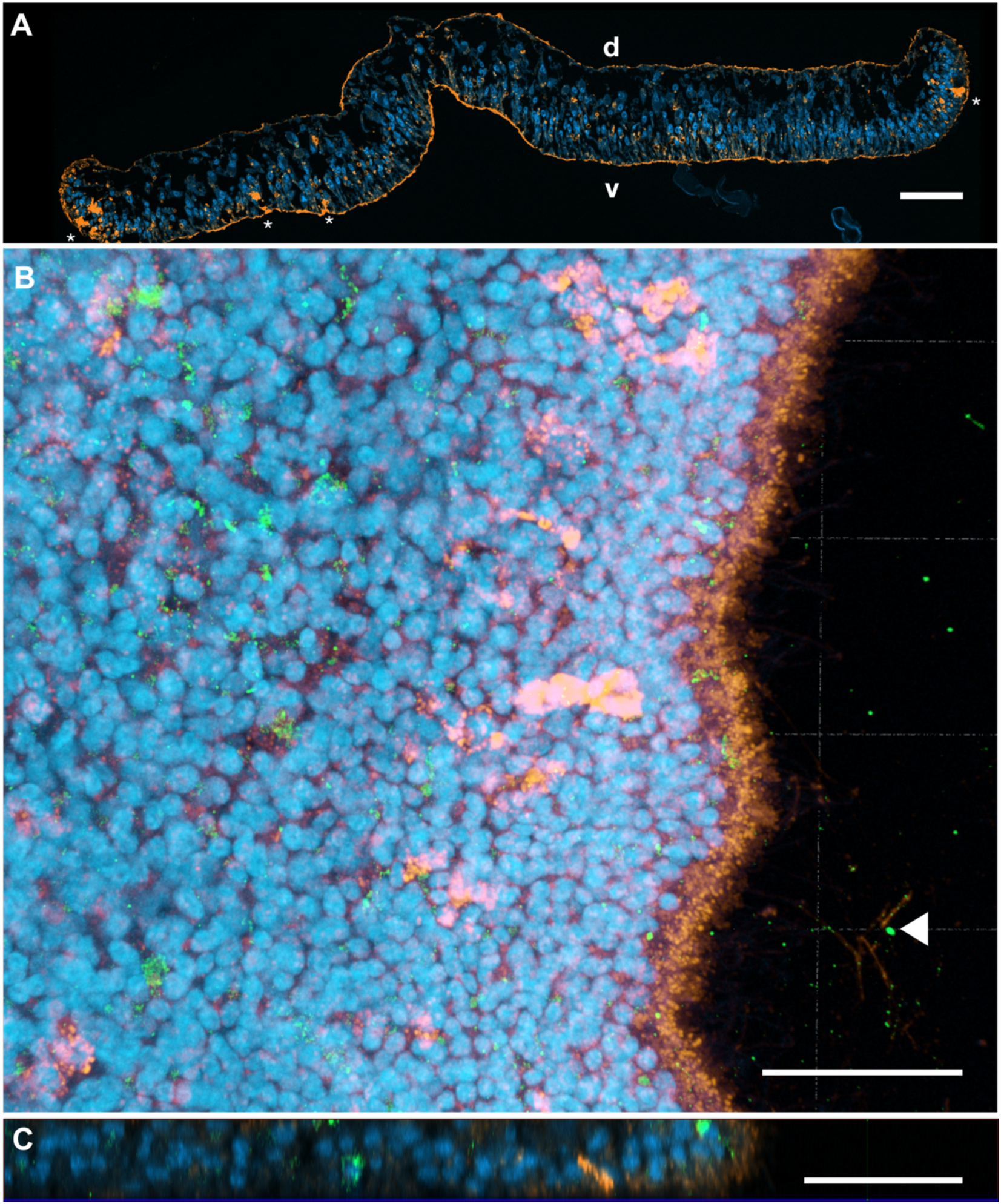
PLys is abundant in gland cells of the ventral epithelium and detectable in secreted mucus. **A)** Cross-section of a resin-embedded *Trichoplax* sp. H2 individual, highlighting dorsal (d) and ventral (v) epithelia as well as mucus-secreting gland cells (asterisks). DAPI = blue, Wheat-germ agglutinin (WGA) = orange. **B)** Dorsal to ventral view on a 3D-projection of a Z-stack of a *Trichoplax*. sp. H2 wholemount stained with anti-PLys (green), WGA (orange) and DAPI (blue). Co-labeling of mucus and PLys in trails of the animal is indicated with an arrowhead. **C**) Partial 2D-orthographic projection (YZ) of the Z-stack shown in Fig. 2B. Scale bars correspond to 20 μm.

### PLys proteoforms show highest lysozyme activity under acidic conditions

Given the contrasting domain architecture of the two forms of PLys, we wanted to understand the role of the cleaved N-terminal domain that characterizes pPLys. We therefore heterologously expressed both pPLys and an mPLys variant recombinantly, where mPLys consisted of the g-type lysozyme domain starting at amino position 112 (M112) of pPLys (mPLys_M112). A comparison of pPLys and mPLys_M112 lysozyme activity at room temperature against freeze-dried *Micrococcus lysodeikticus* revealed high specific activities for both variants compared to hen egg white lysozyme (HEWL) at pH 5.0 (21°C; pPLys = 9 U/pmol; mPLys_M112 = 11 U/pmol; HEWL = 0.5 U/pmol) (Fig. 3B). Both variants had maximum activity under low ionic strength (<50 mM) and at a pH between 5.0 and 5.5 (Fig. 3C-D). The specific activity of mPLys_M112 at its pH optimum (pH 5.5) was three times higher than that of pPLys at its respective optimum (pH 5.0) (Fig. 3E). We also tested the bacteriolytic activity of both proteoforms against viable *Bacillus subtilis* and *Escherichia coli* D31 by assessing the dose-dependent increase in fluorescence of a membrane-impermeable DNA-binding dye (Fig. 3F-G). Both PLys variants showed high activity against *B. subtilis* with LD50 values of 100 nM (mPLys_M112) and 130 nM (pPLys). However, we found no lytic activity against the tested gram-negative *E. coli* strain. The lack of activity against gram-negative bacteria is contrary to the results of PLys by Bam and Nakano (5) but consistent with most metazoan lysozymes being inactive towards gram-negative bacteria (2).

**Figure 3:**
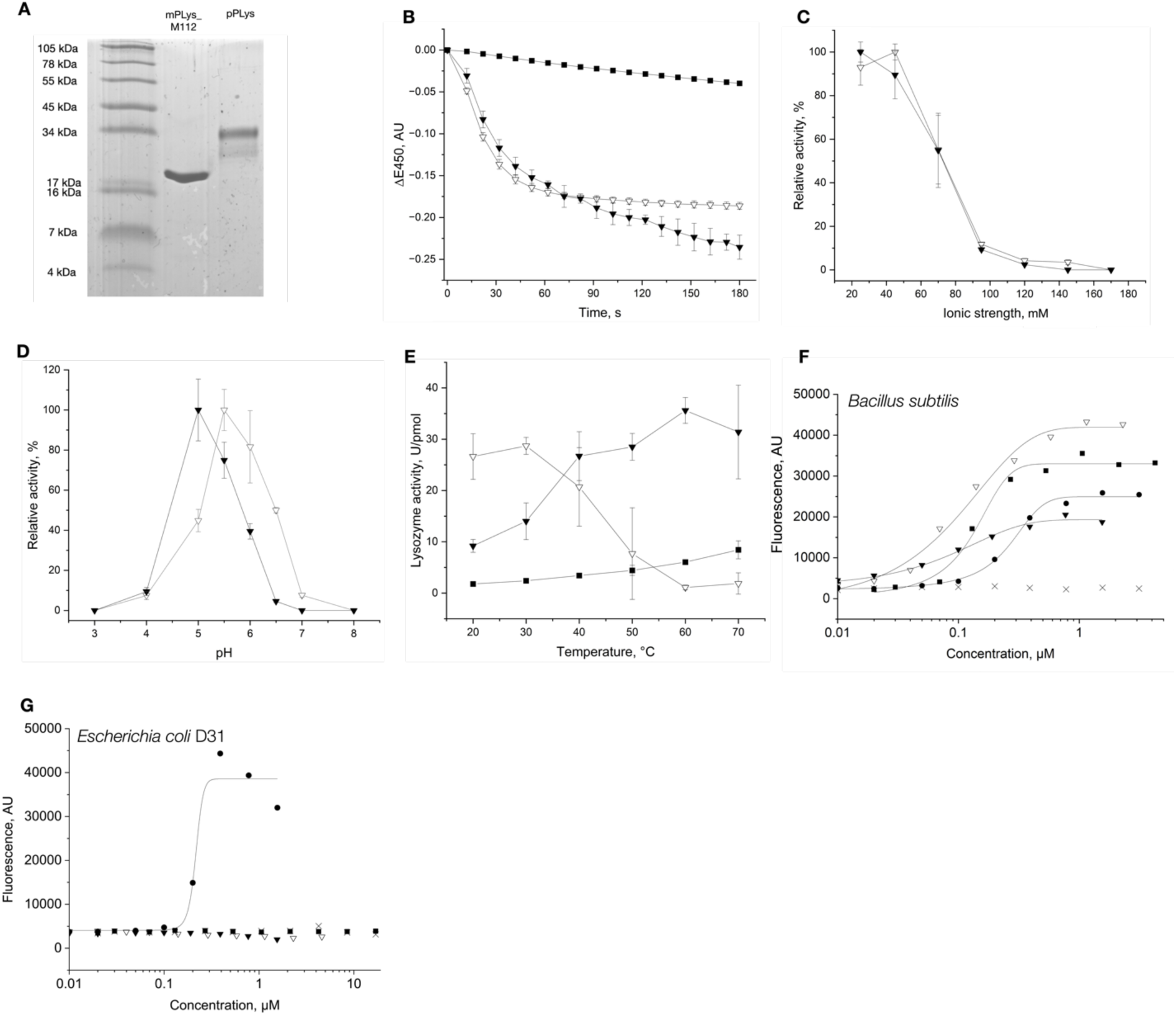
PLys is a potent lysozyme under acidic conditions. **A**): SDS-PAGE gel of purified recombinant mPLys_M112 and pPLys. The molecular mass standard is indicated on the left and marked with the associated molecular masses in kDa. **B**): Lysozyme assay of 32 pmol pPLys, 32 pmol mPLys_M112, and 32 pmol HEWL at pH 5.0. Reaction buffer consisted of 20 mM sodium acetate, pH 5.0 (I = 13 mM). **C,D**): Dependency of lysozyme activity of pPLys and mPLys_M112 on ionic strength (C) and pH (D) at room temperature. **E**): Lysozyme activity of pPLys, mPLys_M112 and HEWL at different temperatures. All measurements were performed in triplicates. **F,G**): Bacterial lysis assay against *B. subtilis* (F) and *E. coli* D31 (G) using a membrane-impermeable dye (SYTOX Green). Fluorescence indicates lysis of cells. Sigmoidal functions were fitted to the dose response curves. mPLys_M112: open triangles, pPLys: closed triangles, HEWL: closed squares, Melittin: closed circles, buffer control: crosses.

### pPLys has a much higher optimal temperature and is active at a broader range of temperatures than mPLys

Since the N-terminal domain of PLys is stabilized by six disulphide bonds, we hypothesized that enzyme activity of the proteoforms could strongly deviate under temperature increase. Our screens across a broad temperature range indeed revealed that the specific lysozyme activity of pPLys increased until 60 °C and virtually stayed at this level at 70 °C, while mPLys_M112 activity peaked at 30 °C and decreased proportional to the increase in temperature (Fig. 3E). Following the strong difference in heat tolerance, we assessed the denaturing dynamics (melting temperatures) for both variants. mPLys_M112 denatured between 25 °C and 35 °C, whereas pPLys denatured between 50 °C and 60 °C, consistent with their temperature-activity profiles (Fig. S4). Overall, the pPLys N-terminal domain appears to have a strong stabilizing effect under physiologically relevant temperatures, at the cost of activity. G-type lysozymes typically contain two disulphide bonds, whereas the PLys lysozyme domain only contains one (18). The increased structural flexibility introduced by the loss of a disulphide bond could explain both increased activity at low temperatures due to an induced-fit type of substrate binding and decreased stability at higher temperatures (19). In pPLys, the cysteine-rich N-terminal domain may compensate for the loss of the disulphide bond and stabilize the protein until maturation to prevent a fast physiological turn-over. We hypothesize that *Trichoplax* uses this trade-off between stability and activity by cleaving the N-terminus from pPLys before secretion. The cysteine-rich repeats (CRRs) which constitute the N-terminus are widely distributed across eukaryotes and were linked to horizontal gene transfer (HGT) events from prokaryotes (20,21). These CRRs are abundant in proteins associated with antimicrobial activity, in which one or multiple CRRs constitute the N-terminus Temperature-dependent activity studies on entire and truncated variants of other proteins with this N-terminal CRR domain structure may elucidate whether the stabilizing effect promotes the selection of this domain architecture and hence has been conserved.

### Trichoplax sp. H2 acidifies its temporary feeding grooves for extracellular digestion

Placozoans form a temporary gastric cavity (feeding groove) only when moving onto their feeding substrate (22). Lipophilic cells and gland cells in the ventral epithelium subsequently show a secretory burst of vesicles, which releases digestive enzymes. In a preliminary study, Mayorova et al. investigated the same ‘gland cell’ annotated meta-cells that are the main expressing cells of *plys* in *Trichoplax* sp. H2. For trypsin, which is also predominantly produced by this cell type, they showed extracellular activity in feeding grooves within digestive events (23). This finding supports our localization of PLys to secretory gland cells and implies that PLys also acts in temporary feeding grooves. Due to the strong dependency of PLys activity on acidic conditions, we hypothesized that placozoans can acidify their feeding grooves despite their temporary nature. To assess the pH during extracellular digestion events, we offered *Trichoplax* sp. H2 zymosan-particles (inactivated yeasts) coupled to a pH-sensitive fluorescent dye (Fig. S5-6) and recorded digestion events in time series (Fig. 4A). While the pH in the feeding grooves initially corresponded to the pH of the surrounding sea water (pH 8.5), we observed spontaneous drops of pH. The pH stayed acidic for multiple tens of seconds and slowly equilibrated back to the initial pH (Fig. S7-8). Over multiple replications of the experiment, we observed an average minimum pH of 6.0 (Fig. 4B), which is close to the activity optimum of mPLys (Fig. 3D). Previous studies on the feeding behavior of *Trichoplax* sp. indicated gland cells and lipophilic cells as primary secretory cells in the feeding process, in which gland cells largely contribute digestive enzymes and lipophilic cells may contribute to the acidification with a secretory burst of vesicles stainable with acidophilic dyes (15). Release of the acidic content of these vesicles in vast amounts may facilitate acidification of the gastric cavity, potentially down to the physiologically optimal conditions for the activity of PLys and other digestive enzymes. Further enzymological characterization of other digestive enzymes of *Trichoplax* spp. may elucidate whether all extracellular digestive processes in placozoans are adapted to acidic conditions.

**Figure 4:**
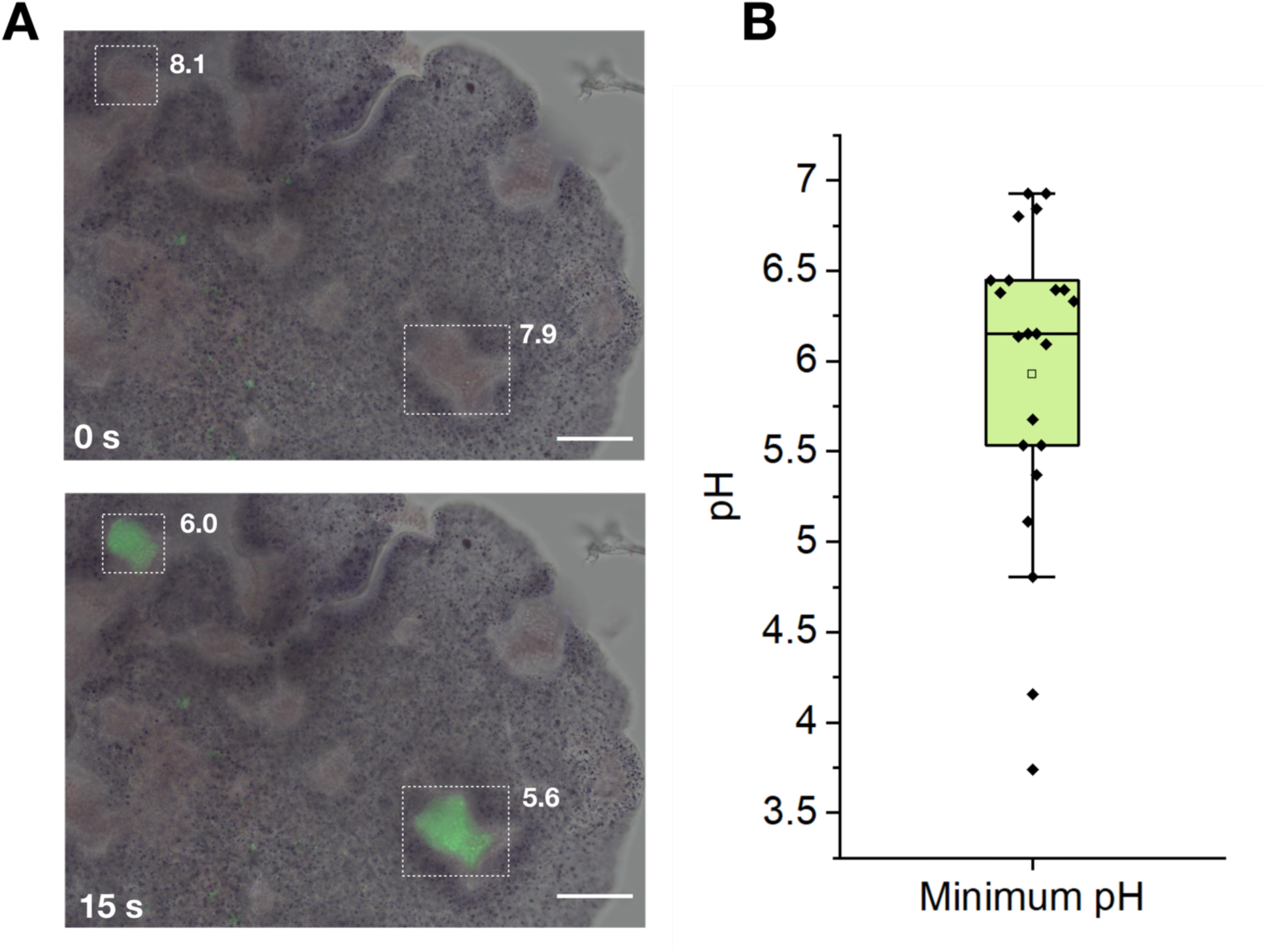
Placozoa acidify their temporary feeding groove pulsatively. **A**): Merged micrographs in brightfield and green-epifluorescence mode of *Trichoplax* sp. H2 feeding on zymosan particles coated with a pH-sensitive dye after 0 s and 15 s. Two feeding grooves are highlighted by dotted lines. The corresponding pH is presented next to the grooves. Scale bars: 50 μm. **B**): Box plot of the minimum recorded pH (closed diamonds) of digestive events (n=22). The open square shows the mean, the horizontal line represents the median.

However, how placozoans lower ionic strength to less than 10% of the surrounding seawater medium remains enigmatic. One possible mechanism may involve the mucus, which is secreted by gland cells in the ventral epithelium. Mucins, the main component of metazoan mucus, are complex proteoglycans which often carry negatively charged building blocks such as carboxy groups, sulfate groups, or sialic acid residues (24). It has already been shown that the negatively charged components of mucins can bind sodium ions and their hydrate shells, causing swelling of the mucus and formation of a biogel (25). Similarly, in placozoans, the nascent mucus in the gastric cavity could potentially take up inorganic cation species from the seawater medium, which is dominated by sodium ions, and lead to a decrease of ionic strength in the aqueous phase inside the gastric cavity. Depending on the pH inside the mucus-loaded exocytotic vesicles of the gland cells, carboxy-groups in the mucus may be initially protonated and lead to proton-sodium exchange between mucus and seawater after secretion, which would also cause a decrease of pH in the gastric cavity. Chemical analysis of placozoan mucus would be necessary to characterize the molecular function of mucus in these early branching metazoans.

### G-type lysozymes presumably originated from an ancient lateral gene transfer

Placozoa are one of the earliest branching animal phyla. As they feature the PLys lysozyme of the GH23 family only, this lysozyme family may have been the earliest in metazoan phylogeny. As the GH23 family also comprises bacterial lytic transglycosylases, this is particularly intriguing. Bacterial lytic transglycosylases also act on peptidoglycan but catalyze the formation of 1,6-anhydromuramyl residues through an intramolecular nucleophilic attack instead of hydrolysis. They are housekeeping genes important for the biosynthesis and turnover of the bacterial cell wall (3). To elucidate the evolution of metazoan g-type lysozymes and to investigate the relationship between GH23 lysozymes and bacterial lytic transglycosylases, we hence constructed a database of all GH23_I members including a set of closely related bacterial lytic transglycosylases that possess the specific sequence motif of the GH23_I. Amino acid sequences were obtained from the carbohydrate active enzyme (CAZY) database, BLAST searches with PLys against nr, and the manual addition of a curated group of GH23_I orthologs detected in metazoan genome assemblies without previously annotated g-type lysozymes (26) (Fig. 5 and S9). We also included sequences of myxobacterial GH23 orthologues, which have been proposed as indicators of a horizontal gene transfer (HGT) from myxobacteria to a metazoan ancestor in a parallel study (27).

**Figure 5:**
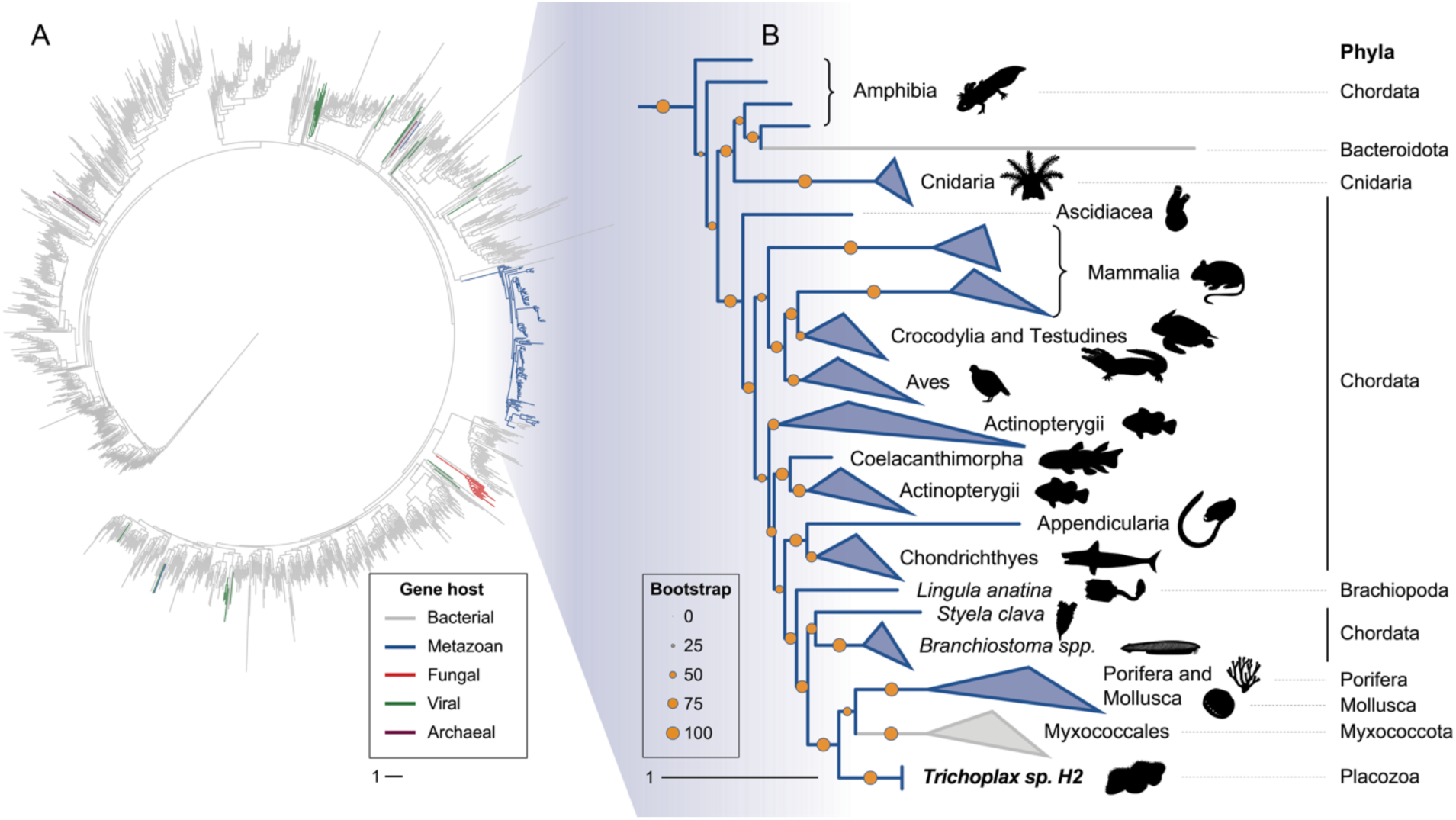
Metazoan g-type lysozymes cluster predominantly into a single clade and are present in six animal phyla. **A**) Phylogenetic tree of 1428 glycoside hydrolase family 23 sequences from all domains of life and **B**) expanded tree branch containing all metazoan sequences. Colors indicate host groups, orange dots show phylogenetic support (aLRT).

In our phylogenetic analysis, out of the 131 metazoan g-type lysozymes included in the dataset, 129 clustered in a single clade (clade support: 99.3), which is nested inside the bacterial lytic transglycosylases. Intriguingly, the manually added myxobacterial orthologues clustered in a monophyletic clade which itself is nested inside the metazoan GH23 lysozyme clade, while other myxobacterial GH23 orthologues annotated as soluble lytic transglycosylases were placed within the bacterial sequence clades. We therefore interpret the myxobacterial sequences clustering in the metazoan clade as professional lysozymes and not lytic transglycosylases, also considering the predatory bacterivorous lifestyle of myxobacteria (28). Additionally, the branch lengths between the myxobacterial lysozymes and the next metazoan relatives are rather short compared to the long branch separating the metazoan clade from the bacterial lytic transglycosylases, which indicates a shorter time frame of molecular evolution. These results indicate the myxobacterial GH23 lysozymes to have originated in a lateral gene transfer of a metazoan GH23 lysozyme to a myxobacterial ancestor, while the metazoan GH23 lysozymes itself originated from a much older HGT of a bacterial lytic transglycosylase to a metazoan ancestor. While the molecular evolution trajectory of most GH23 lysozyme orthologues within the metazoan clade was consistent with the phylogenetic relationships of the respective organisms, we occasionally observed the clustering of orthologues from phylogenetically distant species. This could be due to gene duplication and different evolutionary trajectories of the resulting paralogues or to the horizontal gene transfer between different metazoan ancestors, which has been reported for a subgroup of protostomian g-type lysozymes (27). The prevalence of g-type lysozymes in all early branching animals (Porifera, Cnidaria and Placozoa) except Ctenophora suggests that g-type lysozymes originated during early metazoan evolution. The g-type lysozyme appears to be an innovation of the lineage split from ctenophores and would support the hypothesis of ctenophores being a sister phylum to all other metazoans (29). The gene was then lost in most of the Protostomia except mollusks and brachiopods, while it was retained in most chordate lineages (Fig. 6).

**Figure 6:**
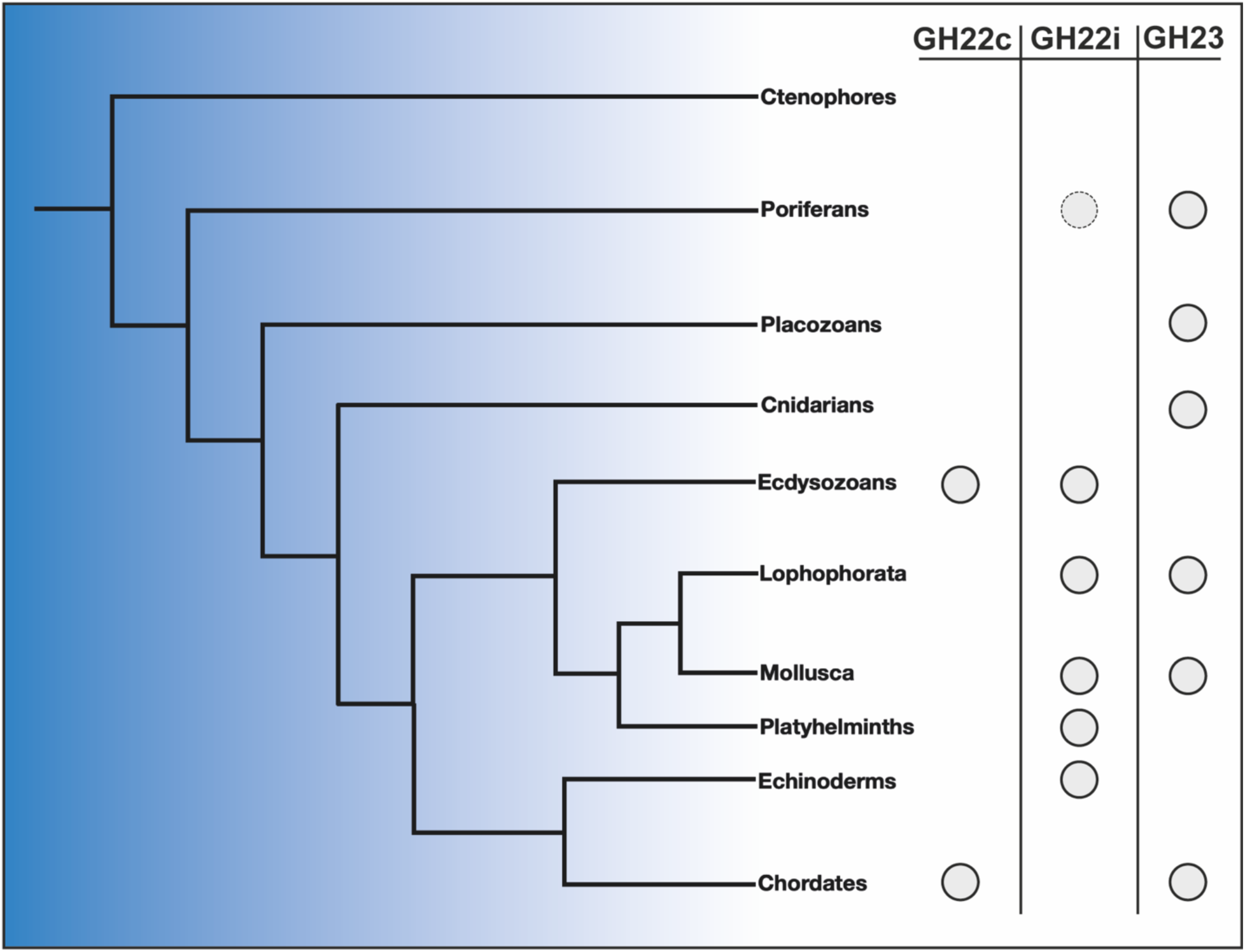
G-type lysozymes are widely distributed over the animal tree of life and the only lysozyme of early all early branching metazoans except ctenophores. Overview phylogram of metazoans according to Laumer et al. (7) that shows the distribution of the most abundant lysozyme families (closed circles). The status of lysozyme abundance is based on Callewaert and Michiels (2), Berndt et al. (30) and this study. As only one GH22i lysozyme has been reported for poriferans (31), the respective circle is dotted.

All early branching animals lack homologues of any other lysozyme subfamily, except a description of an i-type lysozyme in the sponge *Suberites domuncula* (31). In a previous phylogenetic analysis of i-type lysozymes, we showed the *Suberites* lysozyme to be most similar to deuterostomian i-type lysozymes (30). As all other analyzed Porifera lack i-type lysozymes, this could either be a contamination or a secondary acquisition of the i-type lysozyme by an ancestor of *Suberites*. While all metazoan g-type lysozymes cluster together and diverge from bacterial lytic transglycosylases, the exact relationship between these ancient and very similar protein folds, which show low sequence identity but a substantial structural conservation, are difficult to resolve from primary structure analyses. Protein folds and structural sub-motifs of homologous proteins have been shown to be more conserved than their amino acid sequences (32,33). Additionally, the invention of fast and highly accurate protein prediction algorithms allow to generate reliable protein structure models, which can be used for these deep evolutionary analyses (34–36). We therefore used a tertiary-structure based phylogenetics approach to reconstruct deeper relationships within the lysozyme superfamily and included c-type and i-type lysozymes as well as the most closely related GH-families (1449 structures in total) (4,36). The metazoan GH23 lysozyme branch separates from all other metazoan lysozyme families and branches within clades of GH23 bacterial lytic transglycosylases, while the metazoan lysozymes of the GH22i and GH22c subfamilies cluster together with the phage-type lysozymes of the GH24 and GH46 (Fig. 7 and S11).

**Figure 7:**
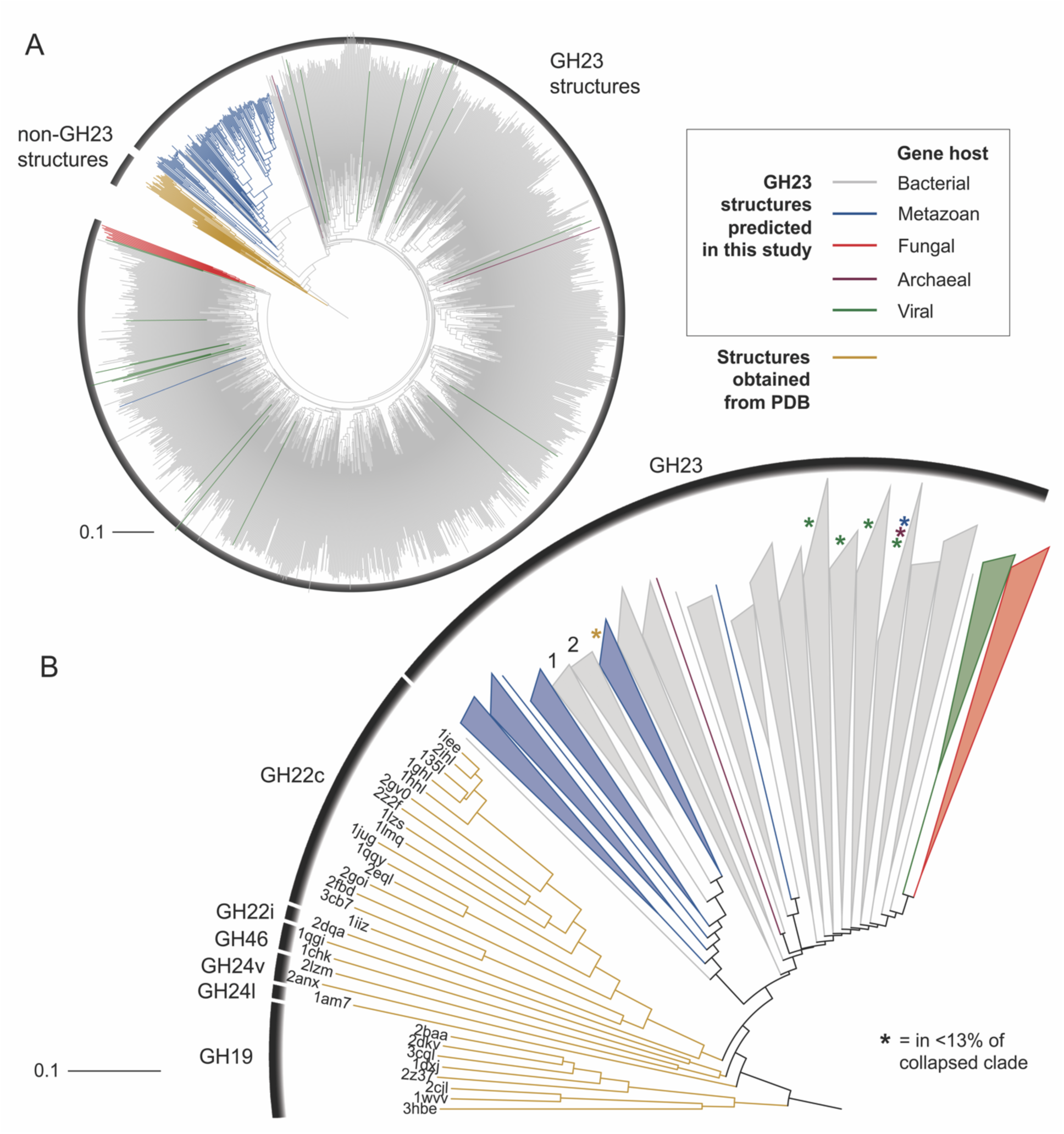
Structure-based phylogeny of GH families within the lysozyme superfamily. In **A**), the whole tree is depicted, in **B**), GH23 clades were collapsed and colored using the same color code as in A). Two clades of myxobacterial lysozyme clades within the metazoan GH23 clade are labeled with “1” and “2”.

This phylogenetic pattern is further strengthened by previous structural biology studies, which showed GH23 enzymes to be more similar to the putative common molecular ancestor of chitinases and murein-active glycosidases than c-type lysozymes (GH22c) and i-type lysozymes (GH22i) in terms of active center organization, reaction mechanism, and fold (4,37). As our sequence- and structure-based phylogenetic analyses indicate, a bacterial lytic transglycosylase may have been horizontally transferred to a metazoan or pre-metazoan ancestor in an ancient event and evolved into the metazoan g-type lysozymes, from where a myxobacterial ancestor acquired a professional GH23 lysozyme gene. To reconstruct the evolutionary trajectories of lysozymes in even higher detail, future genomic analyses of early-branching metazoans and metazoan-related holozoans may allow to provide more comprehensive databases and to identify novel glycoside hydrolase orthologues. Moreover, detailed studies on the abundance and mechanisms of horizontal-gene transfer events involving eukaryotes and prokaryotes are certainly needed to account for the impact of horizontal gene transfer on metazoan evolution.

## Materials and Methods

### *Trichoplax* sp. H2 culture and collection

Organisms of *Trichoplax* sp. H2 strain “Panama” were cultivated at room temperature in petri dishes in 34 PSU filtered sea water (FSW) and fed with *Rhodomonas* sp. Water was changed every two weeks and every culture was discontinued after six weeks. For protein extraction, individuals were picked and transferred into clean reaction tubes with FSW. After transfer of 50 to 100 individuals, the tube was shortly centrifuged to sediment the animals at the bottom. Following the removal of the supernatant, the pellets were stored at -20 °C until use.

### *Trichoplax* sp. H2 lysis and protein detection

Sample pellets comprising 500 individuals of *Trichoplax* sp. H2 in total were lysed using a total volume of 500 μL of 8 M urea, 50 mM sodium bicarbonate buffer, 300 mM NaCl, pH 7.4. The pellets were suspended and pipetted up and down until the lysate appeared homogenous. The lysate was centrifuged at 15,000 x *g* at 4 °C for 15 min to remove insoluble debris. The supernatant was diluted with 50 mM sodium bicarbonate buffer, 300 mM NaCl, pH 7.4 to a final urea concentration of 1.5 M. Subsequently, the sample was subjected to reversed phase fast protein liquid chromatography (FPLC) using a GE ÄKTA Purifier 100 FPLC system (Amersham Biosciences). The sample was loaded on a Hamilton PRP-3 microbore column and subsequently eluted in a linear gradient from 0.1% trifluoroacetic acid (TFA) to 80% acetonitrile (ACN) in 0.1% TFA in 20 column volumes (CV). The eluate was collected in fractions of 1 mL by an autosampler. The fractions were split in two aliquots each and lyophilized. One aliquot of every fraction was subjected to a Western blot analysis.

For protein separation using Tricine-SDS-PAGE, 50 μg of extracted protein were loaded onto a polyacrylamide gel consisting of a 5% stacking gel and a 13% separation gel (38). The SeeBlue Plus2 protein standard (Invitrogen) was used as molecular mass reference. After electrophoresis, proteins were electroblotted onto a PVDF membrane following a standard semi-dry blotting protocol. Successful blotting was verified by staining with Ponceau S. The membrane was washed with Tris-buffered saline (TBS) and TBS with 0.05% Tween 20 (TBST) and afterwards blocked using 1% bovine serum albumin (BSA) in TBST. The membrane was afterwards incubated overnight with a custom polyclonal antibody mix (0.88 μg/mL) against PLys, generated in two rabbits (Eurogentec). One rabbit was immunized with the peptide C-TKEQQMQGGVAAYNF (Res. 239-253), the other one with the peptide C-PYSSTHISQATNILI (Res. 211-225) corresponding to two putatively surface-exposed alpha helices in the lysozyme domain of PLys following protein structure prediction. Antibodies were enriched and purified using affinity chromatography after final bleeding of the animals. After incubation with the primary antibodies, the membrane was washed three times with TBST for 15 min each and incubated afterwards with a monoclonal antibody against rabbit IgG coupled to horseradish peroxidase (HRP) (BioCat) for 1 h. Following three washes with TBST for 15 min, the membrane was developed using the Amersham ECL Advanced Western Blotting Detection kit (GE Healthcare) and a luminescence detector (GE Healthcare) with an exposure time of 5 s. Based on the results of the Western blot, the remaining aliquots of fractions containing putative proteoforms of PLys were subjected to mass spectrometry.

### Top-down proteomics

Lyophilized fractions from the reversed-phase FPLC fractionation (A9, A10) were dissolved in 50 µL 3% acetonitrile (ACN), 0.1% trifluoroacetic acid (TFA) and centrifuged at 20,000 g for 25 min at 4°C. 10 µL of the supernatant was either reduced by 5 mM TCEP for 1 h at 25°C, or analyzed in its non-reduced form by LC-MS. Chromatographic separation was performed on a Dionex U3000 nanoHPLC system equipped with an Accucore C4 column (50 cm × 75 μm, 2.6 μm, 150 Å) coupled online to a Fusion Lumos Tribrid mass spectrometer (Thermo Fisher Scientific) equipped with the High-Field-Asymmetric Waveform Ion Mobility Spectrometry (FAIMS) Pro Interface. The eluents used were: eluent A, 0.05% formic acid (FA); eluent B, 80% ACN, 0.04% FA. The separation was performed over a programmed 90-min run. Initial chromatographic conditions were 4% B for 5 min, then increased to 15% B over 2 min followed by a linear gradient from 15% to 50% B over 60 min then 50 to 90% B over 2 min, and 11 min at 90% B. Following this, an inter-run equilibration of the column was achieved by 10 minutes at 4% B. A constant flow rate of 300 nl/min was employed. Wash runs were performed between each sample injection. Full scan MS spectra were acquired (800-1300 m/z, resolution 120,000, 4 microscans) and fragment scans of the target ions were collected at CV of -15 or -10 (data dependent scans of 2, resolution 60,000) via CID activation at NCE 25. Target ions were chosen after a survey scan that identified the proteoform. Raw data were analyzed against a database containing a Uniprot *Trichoplax sp. H2* proteome (13.02.2023) (12,150 sequences), Uniprot *Rhodomonas* sp. proteins (374 sequences), and common contaminants (cRAP). The search was performed on Proteome DiscovererTM 3.0 using a ProSightPD 4.2 search engine with a High/High crawler Node (Max. RT difference 3 min and a precursor m/z tolerance of 0.5). Single proteoform search node with the sequence of the proteoform was performed with a precursor and fragment tolerances of 100 Da and 10 ppm, respectively.

### Bottom-up proteomics (N terminomics)

Each 15 µL of the fractions 9 and 10 in 3% ACN, and 0.1% TFA were diluted to 100 µL with TEAB then reduced with 10 mM DTT for 30 min at 56°C. Samples were alkylated with 55 mM CAA at 25°C for 20 min in the dark. A first clean-up was performed using SP3 beads (1:25 w/w protein to beads ratio) and ethanol (EtOH) added to 80% (v/v) with mixing for 15 min at 25°C to induce protein binding. The supernatant was removed and beads were washed three times with 80% EtOH. Dimethyl labelling (reductive demethylation) of N-termini was performed by resuspending protein-bound beads in 200 mM HEPES (pH 7.0) and then mixing with formaldehyde (30 mM) and sodium cyanoborohydride (15 mM) at 37°C for 2 h. Formaldehyde (30 mM) and sodium cyanoborohydride (15 mM) was added again to each sample and kept mixing at 37°C overnight. To quench the reaction, Tris buffer (pH 6.8) was added to 0.6 M for 5 h at 37°C. A second clean-up was done by adding EtOH to 80% for 15 min at 25°C. Then, the supernatant was removed and beads were washed three times with 80% EtOH.

Beads were spun down to remove any remaining liquid and then reconstituted in 200 mM HEPES (pH 8). Digestion was performed with trypsin (1:50 enzyme to protein ratio) overnight at 37 °C. Supernatant containing peptides was vacuum dried, then resuspended in 3% ACN, 0.1% TFA and analyzed by LC-MS. Chromatographic separation was performed on a Dionex U3000 nanoHPLC system equipped with an Acclaim pepmap100 C18 column (2 μm particle size, 75 μm × 500 mm) coupled online to a mass spectrometer. The same eluents as for the Top-down experiment were used. The second separation was performed over a programmed 122-min run. After isocratic elution at 4% B for 2 min, a linear gradient from 4% to 50% B over 90 min then 50 to 90% B over 5 min, and 10 min at 90% B was utilized. A constant flow rate of 300 nl/min was employed. Wash runs were performed between each sample injection. Data acquisition following separation was performed on a QExactive Plus (Thermo). Full scan MS spectra were acquired (300-1500 m/z, resolution 70,000) and subsequent data-dependent MS/MS scans were collected for the 10 most intense ions (Top10) via HCD activation at NCE 27 (resolution 17,500). Dynamic exclusion (20 s duration) and a lock mass (445.120025) was enabled. Raw data were analyzed against a database containing a Uniprot *Trichoplax sp. H2* proteome (13.02.2023) (12,150 sequences), Uniprot *Rhodomonas* sp. proteins (374 sequences) and common contaminants (cRAP). The search was performed on Proteome DiscovererTM 2.5 using a SequestHT search engine with 10 ppm and 0.02 Da precursor and fragment ions tolerances, respectively. Digestion with Trypsin_R (semi) with a max of two missed cleavages. Oxidation (15.995 Da) of methionine and dimethylation (28.031 Da) and methylation (14.016 Da) at the peptide N-terminus were set as dynamic modifications. Carbamidomethylation (57.02146 Da) on cysteine and dimethylation on lysine was set as a static modification. Strict parsimony criteria have been applied filtering peptides and proteins at 1% false discovery rate estimation.

### Lysoplate assay

For detection of lysozyme activity, 50 *Trichoplax* sp. H2 organisms were lysed with 500 μL of 50 mM MES buffer, 300 mM NaCl, 1% Triton X-100, pH 5.5. Following lysis, the lysate was centrifuged for at 4 °C at 12,000 x *g* for 10 min to remove insoluble debris. Protein concentration was determined using a Micro BCA assay (Thermo Fisher Scientific). The lysate was concentrated using 10 kDa molecular weight cut-off (MWCO) ultrafiltration inlets (Merck) and 10 μL were loaded onto a lysoplate with 20 mM sodium acetate, pH 5.0, into 3 mm wells (39). The raw extract was loaded onto the plate as well. The lysoplate was incubated at room temperature for 24 h. Ten μL of 2 mg/mL hen egg white lysozyme (Sigma-Aldrich) was used as positive control, lysis buffer as negative control.

### Immunofluorescence of *Trichoplax* sp. H2

*Trichoplax* sp. H2 individuals were either cryofixed using an EM ICE high pressure freezer (Leica) as described previously (11) or by plunging cover slips with animals into 4% paraformaldehyde (PFA) in 1.5X PHEM (PIPES, HEPES, EGTA and MgCl2) buffer, pH 7.0 (40). For high pressure freezing, the animals were first transferred into planchets, covered with hexadecane as filler, and subsequently subjected to high pressure freezing. The vitrified samples were subjected to freeze substitution using a freeze substitution (FS) machine (Leica), following an adapted protocol from Mayorova et al. (16). Briefly, samples were freeze substituted in acetone with 0.5% uranyl acetate at -90 °C for 8 h, followed by incubation at -60 °C for 8 h and at -40 °C for 8 h. After washing with ethanol, samples were progressively infiltrated with medium hardness LOWICRYL HM20 resin at -40 °C (33%, 66%, 100%, each for 2 h), followed by 100% resin overnight. Samples were transferred to fresh resin in 0.2 ml PCR tubes and polymerized under indirect UV-radiation at -40 °C for 48 h, followed by direct UV irradiation for 12 h at room temperature in a nail polish lamp. The hardened samples were remounted and sectioned using a Leica UC7 ultramicrotome and a histo diamond knife (Diatome). Transverse 300 nm thin sections were transferred onto water drops on glow-discharged cover slips and heated until the water evaporated. For histochemical labelling, the cover slips were first blocked with 0.5% fish skin gelatine (Sigma-Aldrich) in phosphate-buffered saline (PBS) for 30 min, followed by incubation with Rhodamine-coupled wheat germ agglutinin (WGA), diluted 1:300 in blocking buffer, at room temperature in a wet chamber for three hours. Subsequently, the cover slips were washed with several changes of PBS and incubated in 1 μg/ml DAPI in ddH_2_O, for 10 min. The cover slips were then washed in two quick changes of ddH_2_O. Excessive water was removed with a filter paper and the cover slips were mounted onto an 8-μL drop of Mowiol (Carl Roth). The cover slips were stored for one day at room temperature at a dark place to ensure hardening of Mowiol and subsequently stored at 4 °C until image acquisition. For wholemount-immunofluorescence, animals were transferred onto cover slips in dishes with artificial sea water (ASW) and allowed to settle for three hours. Subsequently, they were plunged into 4% PFA in 1.5X PHEM buffer and fixed at room temperature for one hour. Afterwards, the cover slips were washed by transferring into fresh PHEM buffer and animals were photobleached for one hour using a high-intensity cold white-light LED (Schott, Germany). Afterwards, the samples were blocked using 0.5% fish skin gelatine in PBS with 0.1% Tween-20 (PBST) for 30 minutes in a wet chamber at room temperature, followed by an overnight incubation with a custom polyclonal antibody mix (Eurogentec) against PLys diluted 1:10 in blocking buffer. A negative control was incubated in blocking buffer only. The next day, the cover slips were washed five times with blocking buffer and incubated 30 minutes with the secondary antibody (antiRabbit-FITC conjugate) diluted 1:100 and WGA-Rhodamine diluted 1:200 in blocking buffer. Afterwards, the cover slips were washed in five changes of PBST and incubated in 1 μg/mL DAPI for five minutes. Excess liquid was blotted away using a filter paper and the cover slips were mounted on drops of Mowiol on slides. Imaging was conducted with an LSM 900 with Airyscan confocal microscope (Carl Zeiss) using a Plan-Apochromat 40x/1.4 Oil DIC M27 objective and three laser excitation wavelengths (405/488/561). Images were acquired in the Airyscan Multiplex mode with tiling and subsequently Airyscan processed and stitched using the Zeiss ZEN v3.2 software.

### Recombinant protein production and purification

cDNA of mPLys_M112 and pPLys, without a signal peptide predicted using SignalP v5.0, was synthesized and inserted into pET-30-a(+) plasmids, codon optimized for *E. coli* (BioCat) (41). The sequence of mPLys_M112 was generated from Uniprot entry B3RIZ1, while the sequence of pPLys was obtained from Uniprot entry A0A369RX22. The amino acid sequence of the pPLys fusion protein (pPlys-FP) was MHHHHHHSSGLVPRGSGMKETAAAKFERQHMDSPDLGTDDDDKAMKSVILLSAFVAVAFAALNDDCNAGQGSCQYDSQCYTGTPASGLCPYDPSNVKCCPKVGQSCKSSTGNCMLTTHCGGTTYSGYCPGPSSVRCCVSSSGSGGSYPTKYGDFMRINPSGASSATARQDGLSYSGVAASNKLASNDYNRCLRYKSQFQSASSSTQIPVGLITAIASRESRCGGALDSNGYGDHGNGYGLMQVDKRYHSLQGGPYSSTHISQATNILISSINGVANNHRSWTKEQQMQGGVAAYNFGVGNVQSIGGMDIGTTGNDYSNDVVARAQWFHNNGFN. The amino acid sequence of the mPLys_M112-FP was MHHHHHHSSGLVPRGSGMKETAAAKFERQHMDSPDLGTDDDDKAMGRINPSGASSATARQDGLSYSGVAASNKLASNDYNRCLRYKSQFQSASSSTQIPVGLITAIASRESRCGGALDSNGYGDHGNGYGLMQVDKRYHSLQGGPYSSTHISQATNILISSINGVANNHRSWTKEQQMQGGVAAYNFGVSNVQSIGGMDIGTTGNDYSNDVVARAQWFHNNGFN.

Besides the lacking N-terminal domain, mPLys_M112 differed from pPLys only in a single amino acid substitution from glycine to serine in amino acid position 256. As this residue is located inside a loop, oriented towards the solvent, and does not form the conserved substrate binding site or active site in the predicted protein structure, we considered this amino acid substitution not to affect either lysozyme activity or protein stability. A chemically competent *E. coli* Shuffle *LysY* strain, which is engineered for cytoplasmic expression of disulphide-bond containing proteins, was transformed with the respective plasmids and grown in LB medium (50 μg/mL Kanamycin) at 37 °C and 150 rpm overnight. The overnight culture was split into 500 μL aliquots and mixed 1:1 (V/V) with glycerol. The resulting glycerol stocks were stored at -80 °C until further use. Prior to recombinant protein expression, 100 mL LB medium was inoculated with 100 μL glycerol stock and grown over night as described above. At the next day, 1 L TB medium (50 μg/mL kanamycin) was inoculated with overnight culture to an OD600 of 0.1. When the culture reached an OD600 of 0.4 to 0.6, protein expression was induced using 0.4 mM IPTG. For protein overexpression, the culture was incubated at 18 °C and 150 rpm for 16 h. Bacteria were harvested by centrifuging at 14,500 x *g* at 4 °C for 30 min. The resulting pellets were resuspended in 50 mM sodium phosphate buffer, 300 mM NaCl, pH 7.4. Bacterial cells were lysed using an MS-73 sonotrode (Bandelin) following the manufacturer’s instructions. The lysate was cleared by centrifuging at 35,000 *x g* at 4 °C for 30 min. The pellet was resuspended in buffer and the lysing and centrifuging step was repeated. The resulting supernatants were collected and subjected to immobilized metal-ion affinity chromatography using HiTrap Talon Crude columns with a CV of 5 mL (Cytiva) and a peristaltic pump (Pharmacia). After sample loading, the column was washed with 10 CV sample buffer. Overexpressed fusion protein was eluted with sample buffer containing 500 mM imidazole. All further chromatographic purification steps were performed on an ÄKTA Purifier FPLC. The eluted fusion proteins were desalted into a 20 mM Tris-HCl buffer containing 50 mM NaCl, 20 % glycerol (w/V) and 2 mM CaCl_2_, pH 6.8, using a HiPrep 26/10 desalting column (Cytiva). Enterokinase, light chain (New England Biolabs) was added in a ratio of 15 ng Enterokinase per milligram fusion protein. After two days of incubation at room temperature, the cleaved protein of interest (POI) was purified using size exclusion chromatography (SEC). For SEC, a Superdex PG 75 16/600 column (Cytiva) was used. After sample loading, the sample was eluted with a 20 mM MES buffer containing 300 mM NaCl and 20% glycerol, pH 5.5, using a flow rate of 0.75 mL/min. Eluted proteins were collected in 1-mL fractions. The purified proteins were snap frozen in liquid nitrogen and stored at -80 °C until use. Protein concentration was determined measuring the absorbance at 280 nm and applying the calculated extinction coefficient at this wavelength.

### Lysozyme assay

For quantification of lysozyme activity, a turbidimetric assay after Shugar was used (42). Lyophilized *Micrococcus lysodeikticus* ATCC No. 4698 (Sigma-Aldrich) (0.2 mg/mL) were suspended in reaction buffer. For standard measurements a 20 mM sodium acetate buffer with varying pH values was used. Different ionic strengths were adjusted with NaCl. To determine the optimal pH value for lysozyme activity, a modified buffer after Ellis was used, containing 5 mM sodium carbonate, 5 mM 2-amino-2-methylpropan-1,3-diol, 5 mM sodium dihydrogen phosphate and 5 mM citric acid (43). For every measurement, 20 μL of lysozyme solution were added to 730 μL *Micrococcus* suspension. The decrease in extinction at a wavelength of 450 nm was measured photometrically. For measurements at room temperature, an EasySpec UV-Vis spectrophotometer (Safas) was used. Measurements at different temperatures were performed with an UV2600i dual-beam absorption spectrophotometer equipped with a CPS-100 temperature-controlled cuvette holder (Shimadzu). One U of lysozyme activity was defined as a decrease of extinction at 450 nm of 0.001per minute (42).

### Bacterial lysis assay

For assaying the lysing potential of pPLys and mPLys_M112, a membrane-permeabilization assay was used, described in detail by Bruhn et al. (44). *Escherichia coli* K12 D31 YpCr (45) and *Bacillus subtilis* (ATCC 6051) were used as target cells in this assay. 1 • 10^6^ CFU/well *E. coli* K12 D31 and 1 • 10^5^ CFU/well in mid-logarithmic growth phase were added to a dilution series of the protein of interest in a black flat-bottom 96-well plate (Sarstedt), together with the DNA binding dye SYTOX Green (Invitrogen). Melittin (Sigma-Aldrich) and Hen egg white lysozyme (Sigma-Aldrich) were used as positive controls, protein storage buffer as negative control. pPLys and mPLys_M112 dilution series were measured in duplicates. Fluorescence was measured with a Tecan Infinite M200 plate reader (Tecan) measuring emission at 538 nm, excited at 495 nm. If possible, sigmoidal curves were fitted to the data in ORIGIN v2023 using the Boltzmann function.

### Thermal shift assay

Stability of recombinant mPLys_M112 and pPLys in different buffers from the Durham pH Screen (Molecular Dimensions #MD1-101) was assessed by thermal shift assays (46). Four µL of 5000x SYPRO Orange dye (Thermo Fisher) were added to 1 mL of protein solution in 100 mM sodium phosphate, 500 mM NaCl, pH 6.5. Ten µL of this mixture were distributed into a 96-well PCR plate (Sarstedt). Ten µL buffer concentrate was added and the plate sealed with a transparent foil (Carl Roth). Melting curves were recorded in an ABI7300 qPCR thermocycler (Thermo Fisher) through the TAMRA channel. The plate was first equilibrated at 21 °C for 2 min and then the fluorescence was measured for 1 min. Subsequently, the temperature was increased in 1-°C increments (0.25 min equilibration followed by 1 min measurement) up to 96 °C. Curves were inspected visually and then processed using *Excel* 16.69.1 (Microsoft) and *Prism* 9.4.1 (Graphpad) as previously described (46).

### *In situ* pH measurement of extracellular digestive events

For assessment of the pH trajectory during feeding events, multiple *Trichoplax* sp. H2 individuales were transferred to a custom microscopic imaging chamber, filled with pHRodo^TM^-Green Zymosan particles (Thermo Fisher) in 34 PSU artificial sea water (100 μg/mL). Imaging was done on a Zeiss Axio Observer 7 inverted microscope with a Zeiss Colibri 7 LED fluorescence source. The Zymosan particles were calibrated using buffers from pH 3 to 9 (50 mM sodium phosphate, 500 mM NaCl). Quantitative imaging was performed using the green fluorescence channel, 100% fluorescence intensity, 150 ms exposure with a 20x Plan-Apo (NA=0.8) objective. For all pH values, the mean fluorescence intensity of Zymosan particles (n=15 per pH) was assessed using Zen Blue v3.9 (Zeiss) (Fig. S5-6). The same imaging settings were used for the *in situ* measurements. Feeding events were followed in a time series with fluorescence and brightfield micrographs every 15 s. The time series were stopped when the animals interrupted feeding and started moving.

### Phylogenetic analyses of g-type lysozyme amino acid sequences

The evolutionary relationships between PLys and other members of the GH23 subfamily I were inferred through Maximum Likelihood (ML) phylogenetics, using the amino acid alignment of the conserved g-type lysozyme domain. From the CAZY database (47), we obtained the list of all 120,967 GH23 annotated proteins (http://www.cazy.org/IMG/cazy_data/GH23.txt; accessed on the 1st of September 2023), and used the accession numbers to retrieve public sequences from the NCBI protein database. The dataset was filtered to keep sequences with a length between 100 and 1,000 residues and that contained both the motives Glu-Ser and Glu-Leu-Met-Gln that are conserved in GH23 subfamily I. Further downsampling was carried out in two steps, first by randomly selecting one sequence from each host species, and then by selecting one sequence per cluster at a 50% identity threshold using CD-HIT v. 4.8.1 (48). Additional metazoan g-type lysozyme sequences were added through rounds of BLASTp searches against the non-redundant sequences database (nr) to improve the representation of the group. Finally, lysozyme sequences from Porifera and Cnidaria from a g-type lysozyme orthogroup (26) and myxobacterial GH23 lysozyme sequences from a parallel study (27) were manually added to the dataset. A chitosanase sequence from *Bacillus subtilis* was included in the dataset as outgroup (acc. num. QHE14361.1). Multiple sequence alignment was carried out with Muscle v.5.1 (49) and the resulting alignment was trimmed to keep residues only that aligned against the g-type lysozyme conserved domain which was determined for PLys sequence through the InterProScan web application (50). The ML tree of the resulting alignment was calculated with IQ-TREE v.2.0.3 (51), based on the general variable time matrix (VT), a FreeRate heterogeneity model with 10 categories (+R10) as selected by ModelFinder (52) and using the SH-aLRT test with 1,000 bootstrap replicates for branch support. Tree representation was obtained with iTOL (53). To account for the simplification generated by the downsampling steps, all eukaryotic sequences removed from the initial dataset, which represented 0.1% of the complete dataset, were placed in the tree following an Evolutionary Placement Analysis (EPA) approach using MAFFT v.7.520 (54) to add the sequences to the alignment and epa-ng v.0.3.8 (55) to build the final tree.

### Protein structure prediction and structure-based phylogenetics of the GH lysozyme superfamily

For protein structure prediction, we used ColabFold v1.5.5 which combines MMseqs2 with Alphafold 2, as previously described (35). Out of five generated models, we chose the highest ranked model as predicted protein structure. Chimera v1.16 was used for molecular visualization (56). Crystal structures of proteins from all GH families within the lysozyme superfamily, following the selection used by Wohlkönig et al. (4) were obtained from PDB database and included in an unrooted ML tree with the predicted protein structure of PLys. The software FoldTree (36) was used to both generate pairwise alignments between the three-dimensional structures and to derive an ML tree based on a local structural alphabets approach.

## Author contributions

HB, ML and HGV conceptualized the study. HB performed most of the experiments and imaging. ID did most of the bioinformatic analysis visualized the respective data. MA and AT performed the mass spectrometry experiments and analysis. MS and AS assisted with the temperature stability assays and the respective analysis. UR and HB prepared the samples for microscopy. HB, HGV and ML wrote the manuscript. HB designed most of the figures. All authors advised the manuscript and approved the final version.

## Competing interests

The authors declare that they have no competing interests.

## Data availability

All proteomics raw data has been uploaded to the ProteomeXchange consortium(57) via the PRIDE partner repository under the accession PXD056545 (Access for reviewers: Username, Password: GXG3RXXb2SzH). Datasets used for the phylogenetic analyses can be downloaded under https://doi.org/10.5281/zenodo.13889706.

## Acknowledgements

We thank Ina Kraus-Stojanowic and Antonia Stanislaus for their support during the recombinant expression and characterization of pPLys, Heidrun Ließegang for assistance of placozoan culturing and the Central Microscopy (CAU) for their support with HPF&FS, ultramicrotomy (Ulrike Voigt and Tanja Zabel) and for providing the Zeiss LSM900 confocal microscope (the German Research Foundation INST 257/650-1 FUGG). Part of this work was performed at the Multi-User CryoEM Facility at the Centre for Structural Systems Biology, Hamburg, supported by the Universität Hamburg and DFG grant numbers (INST 152/772-1|152/774-1|152/775-1|152/776-1|152/777-1 FUGG). We also thank Tim Laugks for technical assistance with high pressure freezing and Brandon Seah for valuable suggestions on the manuscript.

## Notes

### Competing Interest Statement

The authors have declared no competing interest.

### Summary of Updates

The revised version of this preprint includes novel experimental data and discusses recent studies which were included in the analyses and discussions.

