## Supplementary Material for "An ancient lysozyme in placozoans participates in acidic extracellular digestion"

Contents

Supplemental Figures 1 – 10

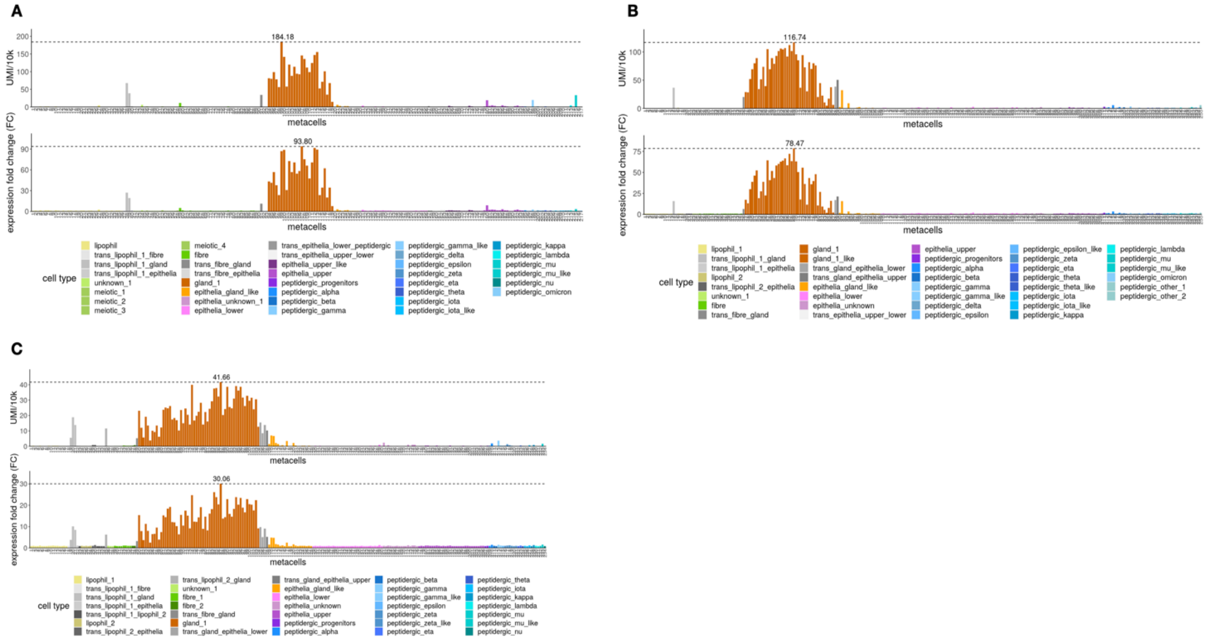


**Supplemental figure 1** Expression pattern of *plys* in A) *Trichoplax* sp. H2, B) *Hoilungia hongkongensis* H13, and C) *Cladtertia collaboinventa* H23. The figure was created using the Placozoa Cell Atlas (Najle et al. 2023).

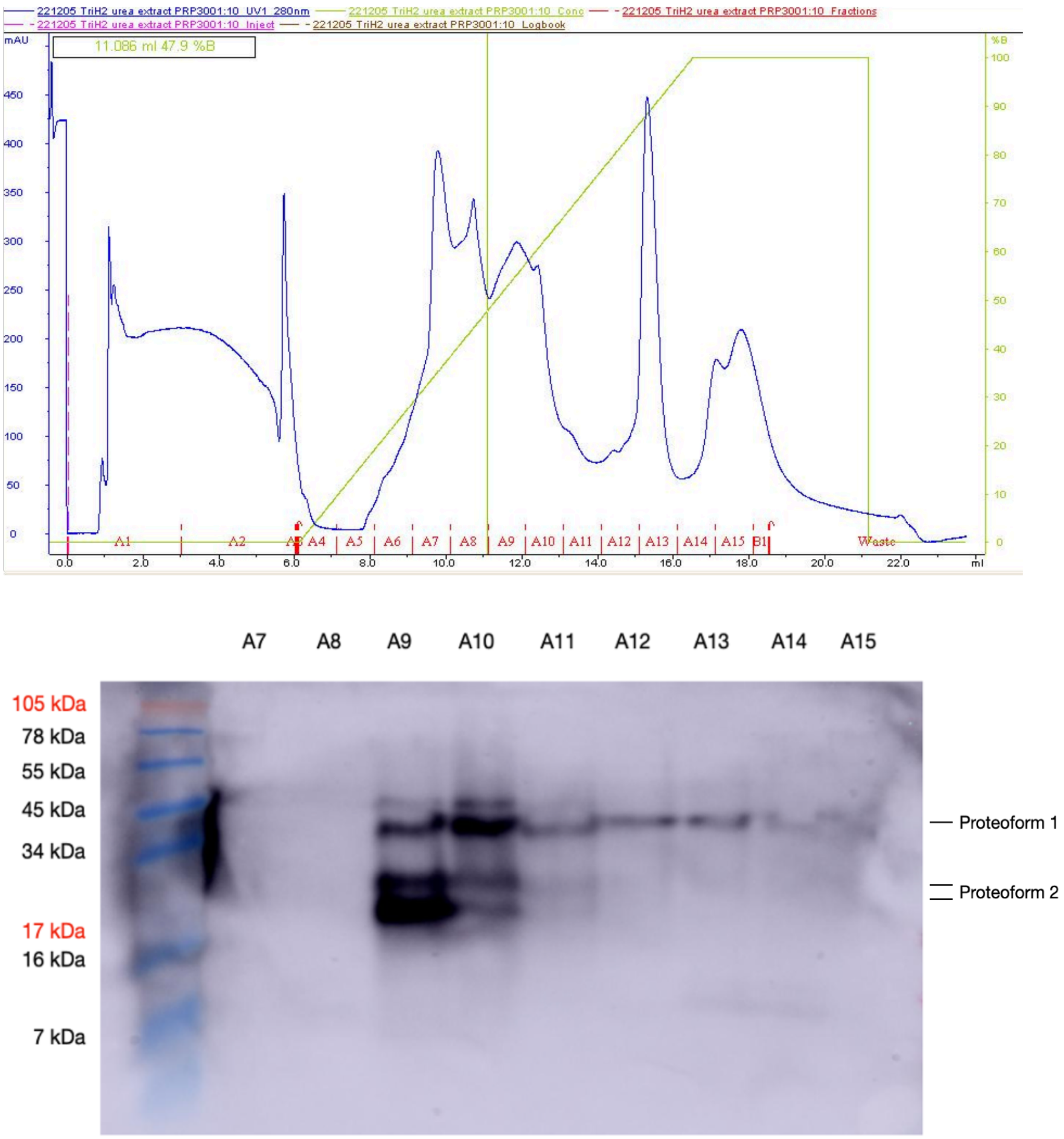


**Supplemental figure 2** Chromatogram of the reversed-phase *Trichoplax* sp. H2 protein extract fractionation and Western blot of fractions A7 to A15 with affinity purified anti-PLys IgG. Invitrogen SeeBlue Plus2 standard was used as molecular mass reference for the Western Blot. The two PLys proteoforms are annotated as evidenced by mass spectrometry.

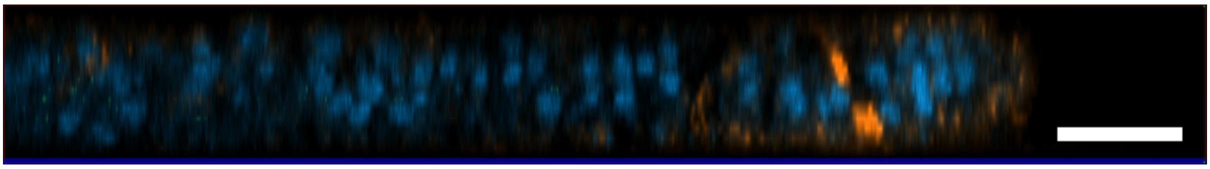

**Supplemental figure 3** Partial 2D-Orthographic projection (YZ) of a Z-stack through a wholemount *Trichoplax* sp. H2 after immunohistochemical labeling with antiRabbit-FITC conjugate, WGA-Rhodamine, and DAPI. Scale bar corresponds to 15 μm.

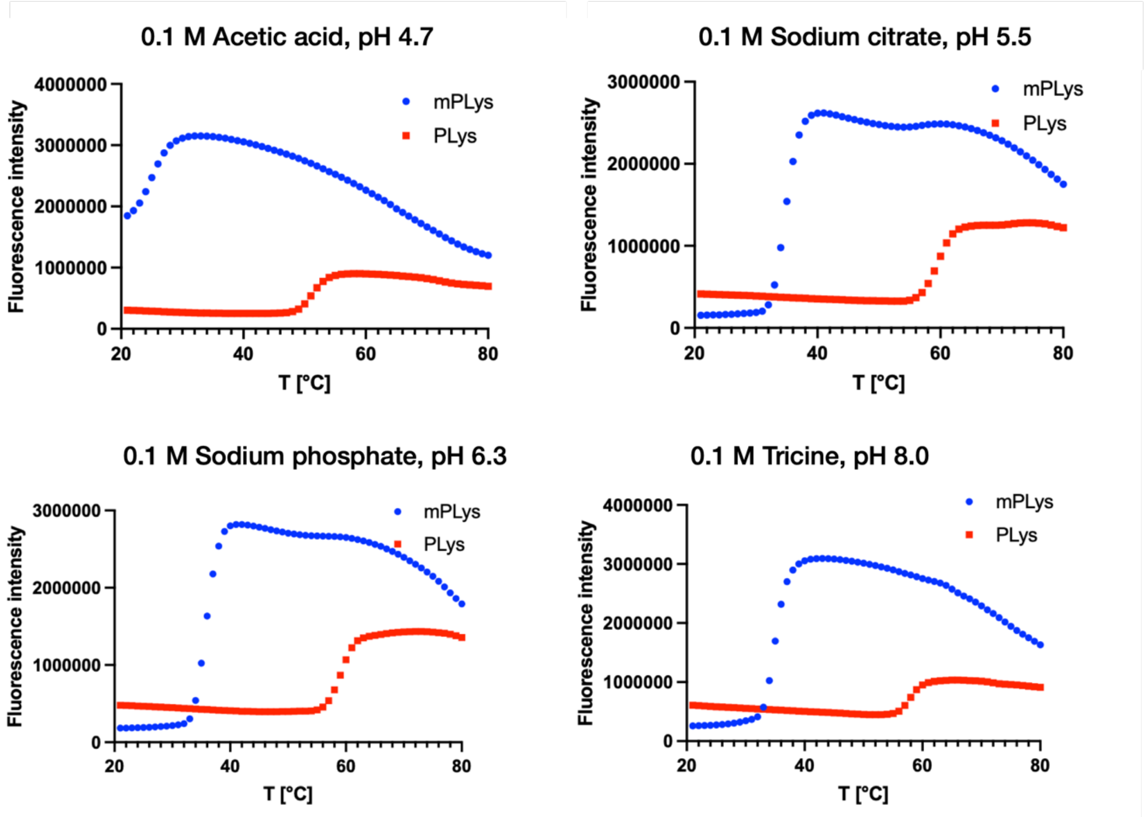


**Supplemental figure 4** Representative melting curves of recombinant mPLys_M112 (blue) and pPLys (red) under different buffer conditions.

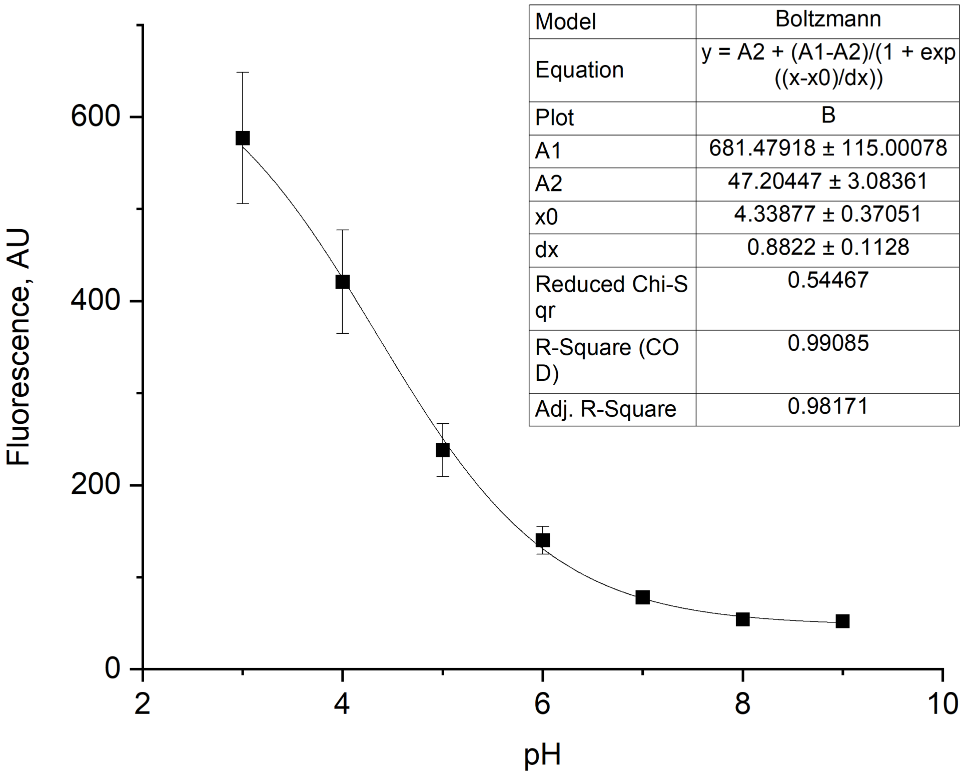


**Supplemental figure 5** Calibration curve of pHRodo™ Green zymosan green fluorescence at differing pH values (n=15 per pH). A sigmoidal curve was fitted to the data.

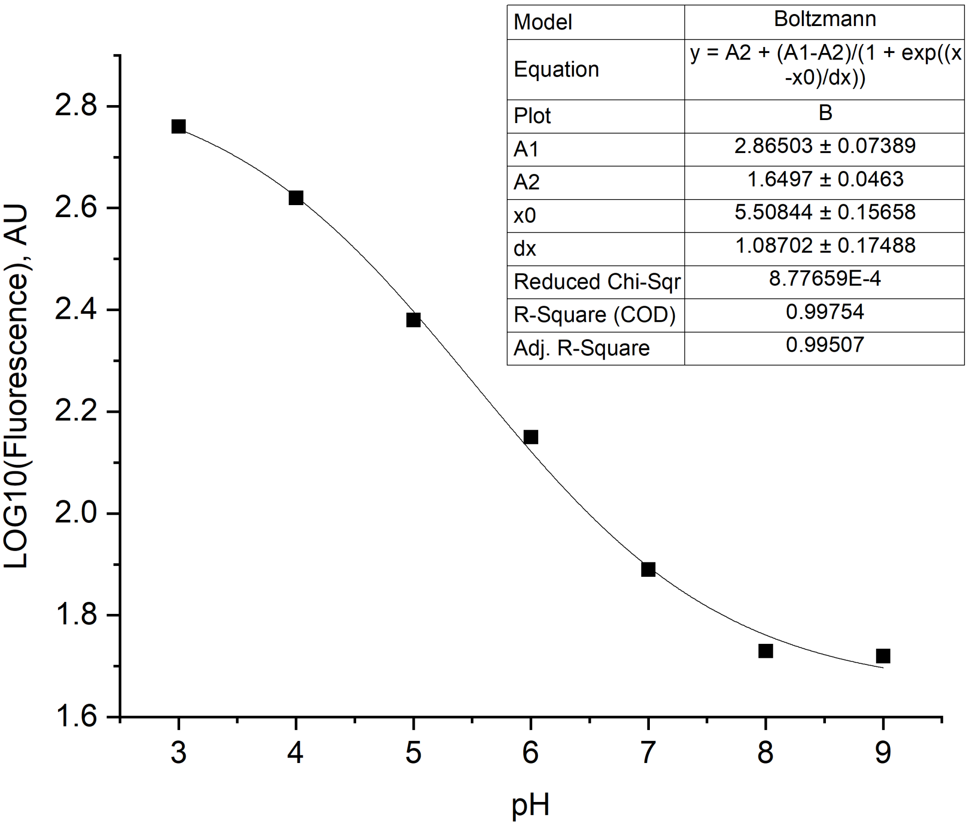


**Supplemental figure 6** Calibration curve of pHRodo™ Green zymosan logarithmized green fluorescence at differing pH values (n=15 per pH). A sigmoidal curve was fitted to the data.

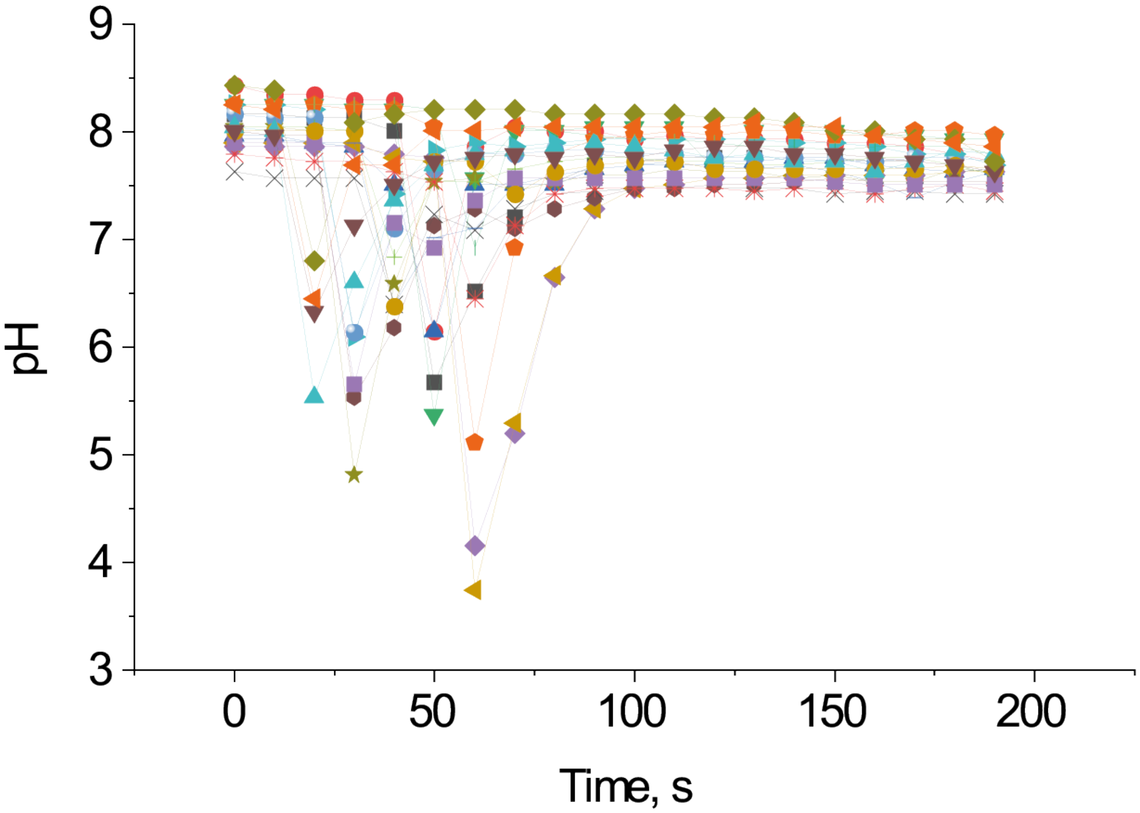


**Supplemental figure 7** pH trajectory during extracellular digestive events (n=22) of *Trichoplax* sp. H2. Each feeding event is represented by an individual symbol.

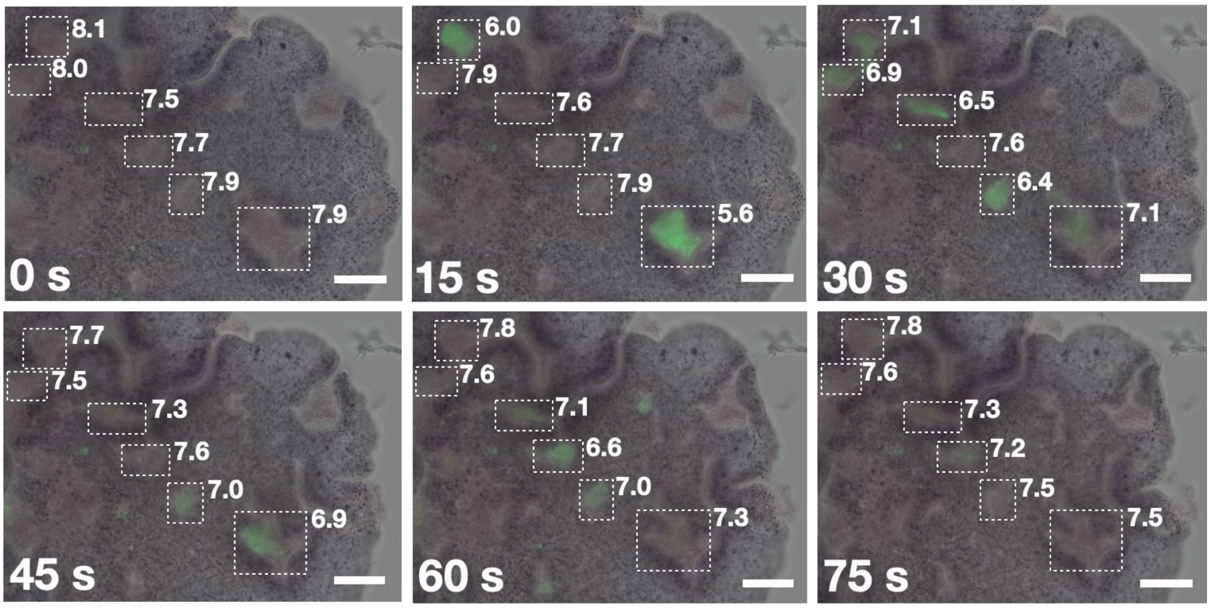


**Supplemental Figure 8** Merged micrographs in brightfield and green-epifluorescence mode of *Trichoplax* sp. H2 feeding on zymosan particles coated with a pH-sensitive dye at different time points. Feeding grooves are highlighted by dotted lines. The corresponding pH is presented next to the grooves. Scale bars: 50 μm.

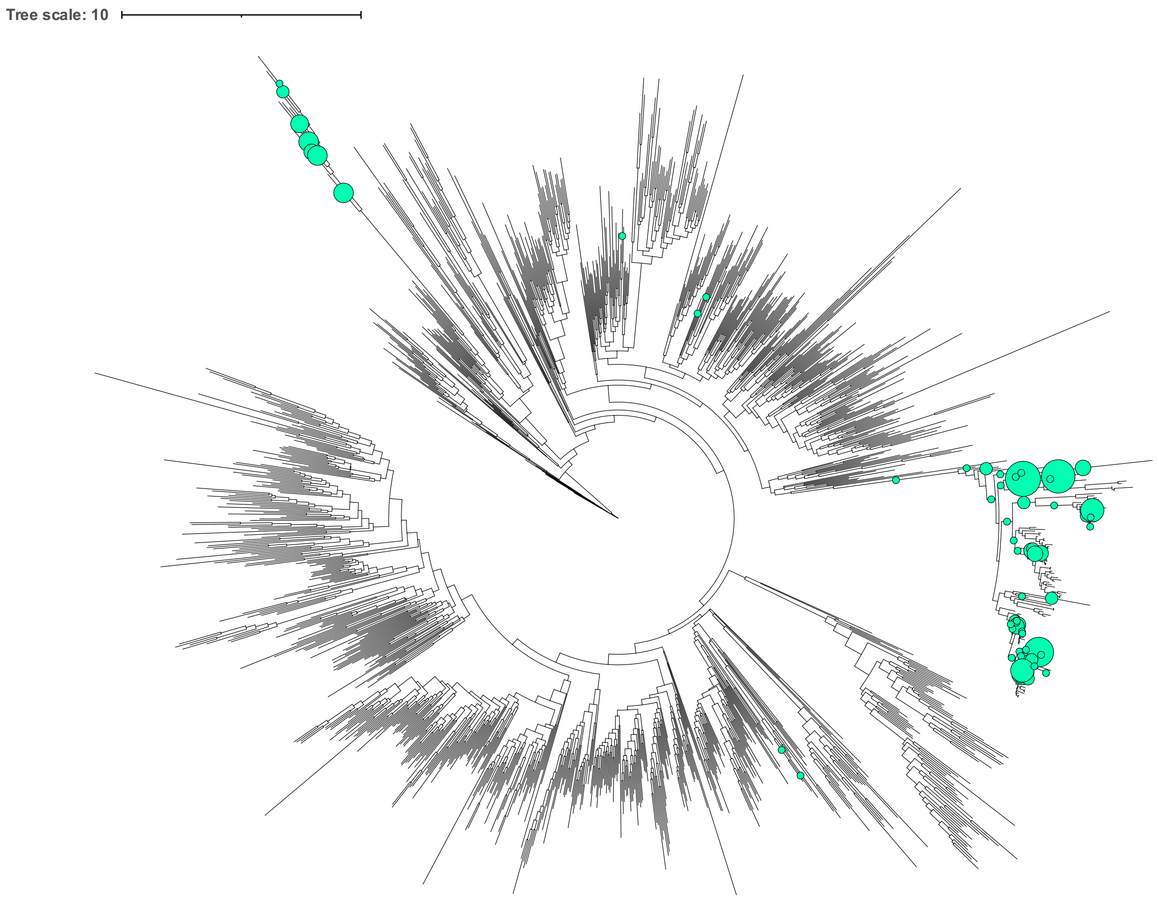


**Supplemental figure 9** Phylogenetic tree of glycoside hydrolase family 23 proteins with evolutionary placement analysis results. Circle sizes are proportional to the number of eukaryotic sequences that were placed into a branch following the EPA.

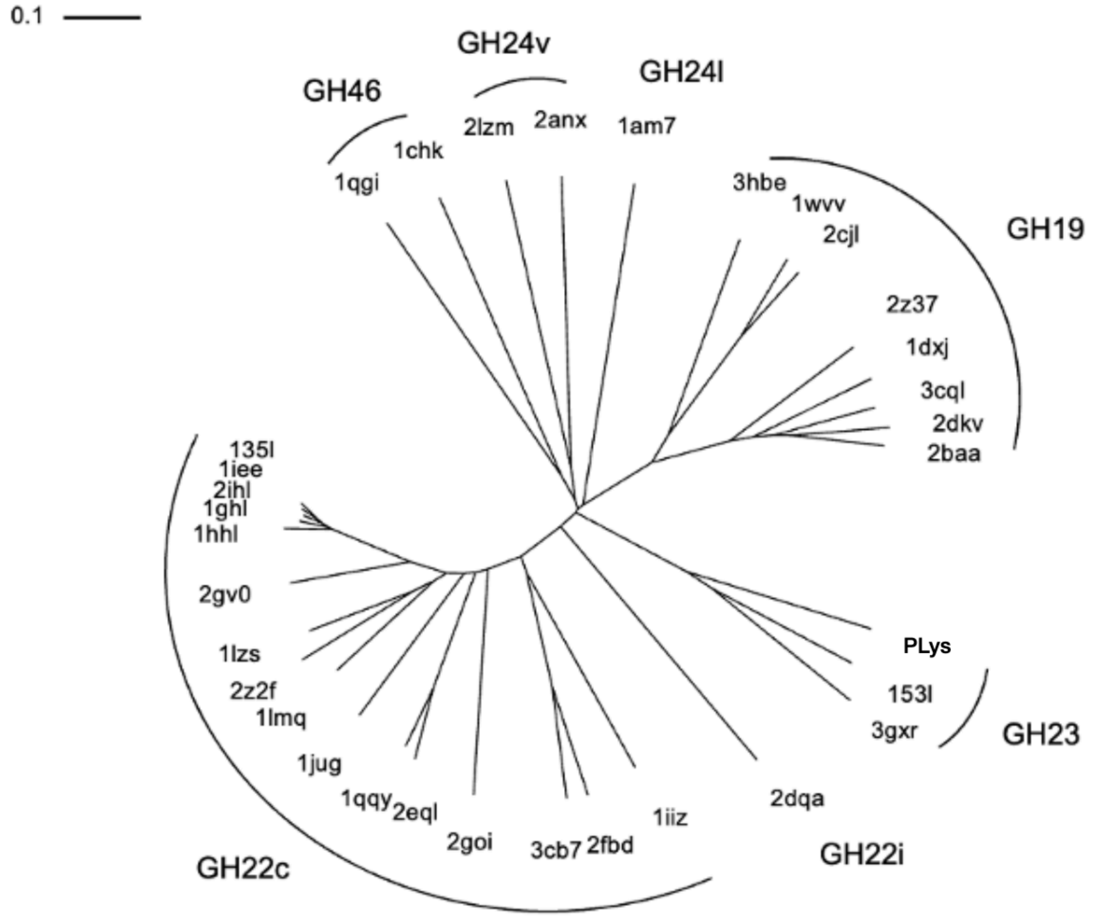


**Supplemental figure 10** Structure-based phylogenetic tree of experimentally solved crystal structures of proteins from the lysozyme superfamily and the model of PLys.
